# Longitudinal Fragility Phenotyping Predicts Lifespan and Age-Associated Morbidity in C57BL/6 and Diversity Outbred Mice

**DOI:** 10.1101/2024.02.06.579096

**Authors:** Alison Luciano, Laura Robinson, Gaven Garland, Bonnie Lyons, Ron Korstanje, Andrea Di Francesco, Gary A. Churchill

## Abstract

Aging studies in mammalian models often depend on natural lifespan data as a primary outcome. Tools for lifespan prediction could accelerate these studies and reduce the need for veterinary intervention. Here, we leveraged large-scale longitudinal frailty and lifespan data on two genetically distinct mouse cohorts to evaluate noninvasive strategies to predict life expectancy in mice. We applied a modified frailty assessment, the Fragility Index, derived from existing frailty indices with additional deficits selected by veterinarians. We developed an ensemble machine learning classifier to predict imminent mortality (95% proportion of life lived [95PLL]). Our algorithm represented improvement over previous predictive criteria but fell short of the level of reliability that would be needed to make advanced prediction of lifespan and thus accelerate lifespan studies. Highly sensitive and specific frailty-based predictive endpoint criteria for aged mice remain elusive. While frailty-based prediction falls short as a surrogate for lifespan, it did demonstrate significant predictive power and as such must contain information that could be used to inform the conclusion of aging experiments. We propose a frailty-based measure of healthspan as an alternative target for aging research and demonstrate that lifespan and healthspan criteria reveal distinct aspects of aging in mice.

## Introduction

Aging studies characterize normative aging, evaluate purported lifespan interventions, and seek to unravel age-specific disease mechanisms. For many aging studies, the primary outcome is natural lifespan. Studies in murine models greatly accelerate these research programs. The typical laboratory mouse lifespan is roughly three years, compared to 80-year human life expectancy [1]. Still, researchers input years of effort to conduct murine aging studies. The ability to ascertain lifespan in murine models without waiting for census events to occur presents an important opportunity to accelerate aging research [2]. One potential approach to reduce the time required for longitudinal studies of aging is to develop predictive tools that reveal an animal’s lifespan antemortem [3]. A second important goal for longitudinal studies of aging is the inclusion of measures of the proportion of life lived in good health (i.e., healthspan) [4, 5]. Although invasive bioassays may be predictive of lifespan (e.g., [3, 6–10], viable proxy endpoints in aging studies would need to be noninvasive to not bias trial outcomes and easy to execute to support frequent assessments and high temporal resolution. Several frailty indices (FI) have been proposed; these are typically comprised of non-invasive and easily observable phenotypes associated with the aging process [11–13].

One approach to lifespan prediction is to determine the onset of late-life decline (LLD), which is a period of increased vulnerability and morbidity before death. LLD can vary widely among individuals and populations, and it is often influenced by environmental and genetic factors [14]. We are aware of only two noninvasive tools to predict life expectancy (LE) in longitudinal aging studies in mice: the temperature x body weight (TxW) measure [15] and the Analysis of Frailty and Death (AFRAID) clock [16]. The TxW composite measure was the first noninvasive computational tool to predict LLD in mice. The rationale behind TxW is based on the observation that body weight and temperature tend to decline in aged mice as they approach natural death. By collecting these routine measures and applying a post-hoc criterion of terminal decline, the TxW developers were able to identify LLD in outbred Hsd:ICR (CD1) mice. They also validated their method in an infectious disease model [17] and three cohorts of aged, inbred strains [18]. They found that TxW can successfully signal the need for more intensive clinical evaluation. However, they noted some limitations, such as the possibility of spontaneous reversal of temperature dysregulation or weight loss in some aged inbred mice, which could lead to premature euthanasia if TxW is used as a predictive endpoint. Therefore, they suggested that other researchers should test their approach in different settings and populations. To our knowledge, no external validation of the TxW measure has been conducted by an independent research group. In this study, we evaluate the performance and applicability of the TxW measure.

Another noninvasive computational tool to measure LLD in mice is the AFRAID clock [16], which uses machine learning (ML) algorithms to derive predictions from the items in a standardized frailty exam. The AFRAID clock is based on the observation that frailty, which is a state of increased vulnerability and reduced resilience to stressors, increases with age and predicts mortality in mammals. By collecting longitudinal frailty data in a cohort of aged male C57BL/6J mice, the AFRAID developers were able to construct a model that estimated the remaining lifespan and the optimal time for euthanasia. They also tested their model in two different interventions that are known to affect longevity: methionine restriction and enalapril treatment. They found that AFRAID predicted an increase in lifespan for methionine-restricted mice but not for enalapril-treated mice. The validity and generalizability of the AFRAID clock were re-evaluated in a retrospective analysis of frailty as defined by Schultz *et al*. [16] in mice under different intervention conditions [19]. They found that AFRAID scores did not correlate with lifespan across the whole cohort and showed inconsistent bias and precision in mortality age estimation across treatment groups, contrary to the original study. They suggested that the AFRAID clock may not be robust or applicable to different settings and populations, and that future studies should focus on building clocks with different frailty features in a variety of interventional cohorts. Therefore, there is a need for further evaluation and improvement of frailty-based ML algorithms as a tool for measuring LLD in mice.

We aimed to further develop predictive endpoint criteria for aging studies that would perform well across genetic background and sex, without regard to specific diagnosis and without differential performance across intervention types. We developed a modified frailty assessment, the Fragility Index (FgI) derived from existing FIs with additional deficits selected by Jackson Lab veterinarians based on their extensive experience in the evaluation and treatment of mice in aging studies (**eTable 1**). We incorporated the FgI within ongoing studies of aging that included mice of both sexes, different genetic backgrounds, and aging interventions to provide data on end-of-life health trajectories and to determine if we could establish generalizable antemortem endpoints that improve clinical decision-making and accelerate longitudinal aging studies.

## Results

The Longitudinal Fragility Study enrolled 246 female Diversity Outbred (DO) mice from a dietary intervention study [20] that was initiated with 960 mice across five diet groups (192 mice per group). All 246 included mice had survived to enrollment age of approximately 2.5 years including 26 (13.5%) in the ad libitum (AL) group, 34 (17.7%) in the 1 day/week fasting (IF1) group, 51 (26.6%) in the 2 day/week fasting (IF2) group, 56 (29.2%) in the 20% calorie restriction (CR20) group, and 79 (41.1%) in the 40% calorie restriction (CR40) group. We also enrolled 43 C57BL/6J (B6) mice (15 female and 28 male) at an age of approximately 2 years. The B6 mice were fed ad libitum. Temperature, body weight, and FgI were assessed weekly across the two cohorts (**Figure 1**).

**Figure 1:**
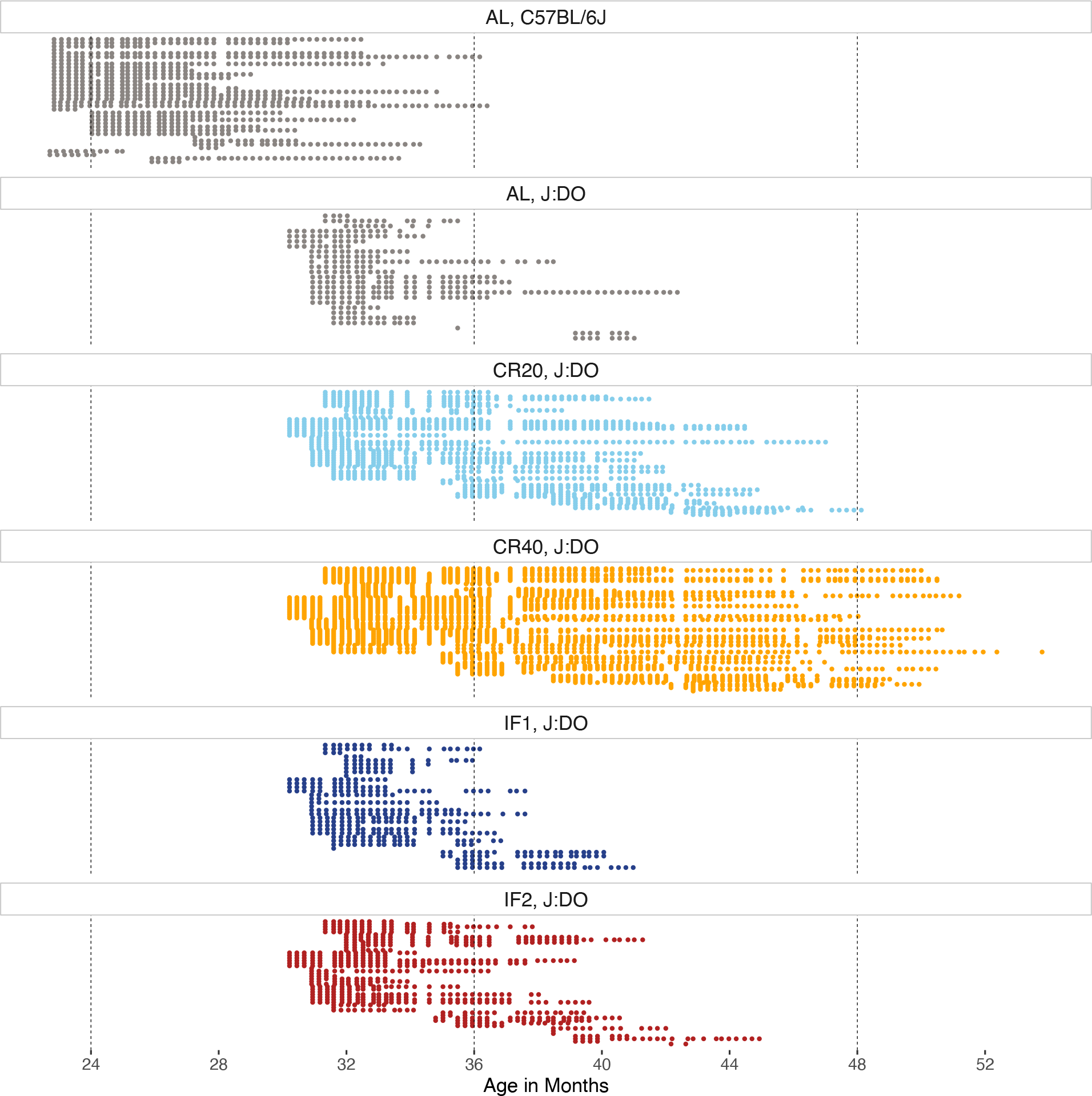
Weekly clinical Fragility Index assessments conducted among aged mice between enrollment and exit resulted in 5,957 total assessments (median [Q1, Q3] = 18 [9, 30] assessments per mouse) collected between September 2019 and December 2021. Each row represents a unique mouse with natural lifespan data (N = 270 total). High temporal resolution achieved across two genetic backgrounds and five experimental conditions. AL = ad lib (grey); IF1 = 1-day intermittent fast (dark blue); IF2 = 2-day intermittent fast (red); CR20 = 20% calorie restriction (light blue); CR40 = 40% calorie restriction (orange).

### Lifespan in the Longitudinal Fragility populations

DO mice were enrolled in the study at 30 months of age (mean=32.7 months). All mice in the DO cohort were females. Median lifespan was 37.4 months and the 90th percentile was 46.3 months. The longest-lived mouse in the DO cohort reached 53.9 months of age (4.5 years). Lifespans were highly variable among DO mice with an interquartile range of 7.2 months (Q1 = 34.3 to Q3 = 41.5 months). We observed significant variation in both mean (p = 3.12e-15) and maximum lifespan (90th percentile, p = 4.1e-08) across diets. AL mice had the shortest median lifespan of 34.6 months, and 40% CR had the longest median lifespan of 40.8 months.

B6 mice of both sexes were enrolled at approximately 24 months of age. Median lifespan was 29.3 months and the 90th percentile was 34.7 months. The longest lived B6 mouse (a male) reached 36.6 months of age. The B6 mouse lifespans were less variable compared to the DO mice, with an IQR of 5.3 months (Q1 = 27.4 to Q3 = 32.7 months). Median lifespan of female mice was 28.9 months (n = 12) and median lifespan of male mice was 29.6 months (n = 24). Neither the mean (p = 0.51) nor the maximum lifespan (p = 0.28) was significantly different between sexes. Male lifespans were more variable compared to females (p = 0.05).

The lifespans reported in these cohorts are conditional on survival to enrollment age and thus are longer than lifespans of unselected mice. The enrollment ages for DO and B6 mice are substantially different which precludes cohort comparisons, except to note that B6 lifespans were less variable compared to the DO lifespans.

### Longitudinal clinical observation identifies normative patterns of aging in mice across sex, diet, and genetic background

We collected 5,957 weekly assessments on mice between September 2019 and December 2021, including two quantitative assays (bodyweight [grams], temperature [Celsius]) and 30 ordinal assays from our Fragility Index (Methods). Median age at the first assessment was 22.8 months for B6 mice, and 31.3 months for DO mice.

FgI scores were obtained as mean level of deficit across all ordinally scored items, with a potential range from 0 to 1, where 1 would indicate maximum health deficit. FgI scores for B6 mice ranged from 0.02 to 0.48, and for DO mice scores ranged from 0.02 to 0.57. The median [IQR] FgI score was 0.22 [0.17, 0.27] and 0.27 [0.2, 0.33] among C57BL/6J and J:DO mice, respectively. Humans also show a sub-maximum threshold of FI scores, although it is typically higher (∼0.7) [21]. The maximum number per-mouse of health deficits rated severe was similar across genetic backgrounds (median [IQR] deficits = 8 [6, 13] among B6 mice and 10 [8, 11] among DO mice).

Several indicators were never observed. For example, no mice presented mild or severe signs of nasal discharge. Cumulative incidence of some indicators was high, indicating that all or nearly all mice eventually showed clinical signs of severe frailty in that health domain. Indicators with high cumulative incidence across both genetic backgrounds included piloerection, kyphosis, and poor grooming (coat condition). Cumulative incidence of individual items was sensitive to diet assignment in the DO mice (**eTable 2**). Mice assigned to the AL group were less likely to experience severe body condition, ocular degeneration, poor grooming, reduced skin turgor, hunched posture, and severe disequilibrium than mice in other diet groups. Two potentially lethal items were more common in the AL group: palpable masses and distended abdomen. One possible interpretation is that dietary interventions increased lifespan but had adverse effects on some aspects of late life health in the DO mice. Survivorship bias is another possible explanation (see below). Comparisons by sex among B6 mice showed few notable differences (**eTable 3**). Activity, head piloerection, and hunched posture deficits were more common among males than females; peri-retro-orbital swelling was rarely observed but marginally more common among the B6 females.

### Age and concurrent FgI correlate with life expectancy differently across genetic backgrounds

To understand the potential for FgI to predict lifespan, we examined how the FgI score changes as a function of the proportion of life lived (PLL) (**Figure 2b**). PLL scaling facilitates comparison of terminal trajectories for mice despite being enrolled at varying ages and allows us to examine characteristics at end of life whether the lifespan was typical (e.g., 2 years) or atypical (e.g., 4 years). For both B6 and DO mice, visual inspection suggested an inflection point in FgI status near 0.95 PLL. This a highly desirable feature for assessment of imminent mortality. Body weight trajectories show similar inflection point in DO mice, but perhaps a broader timescale of decline in B6 males and females (**Figure 2c**).

**Figure 2:**
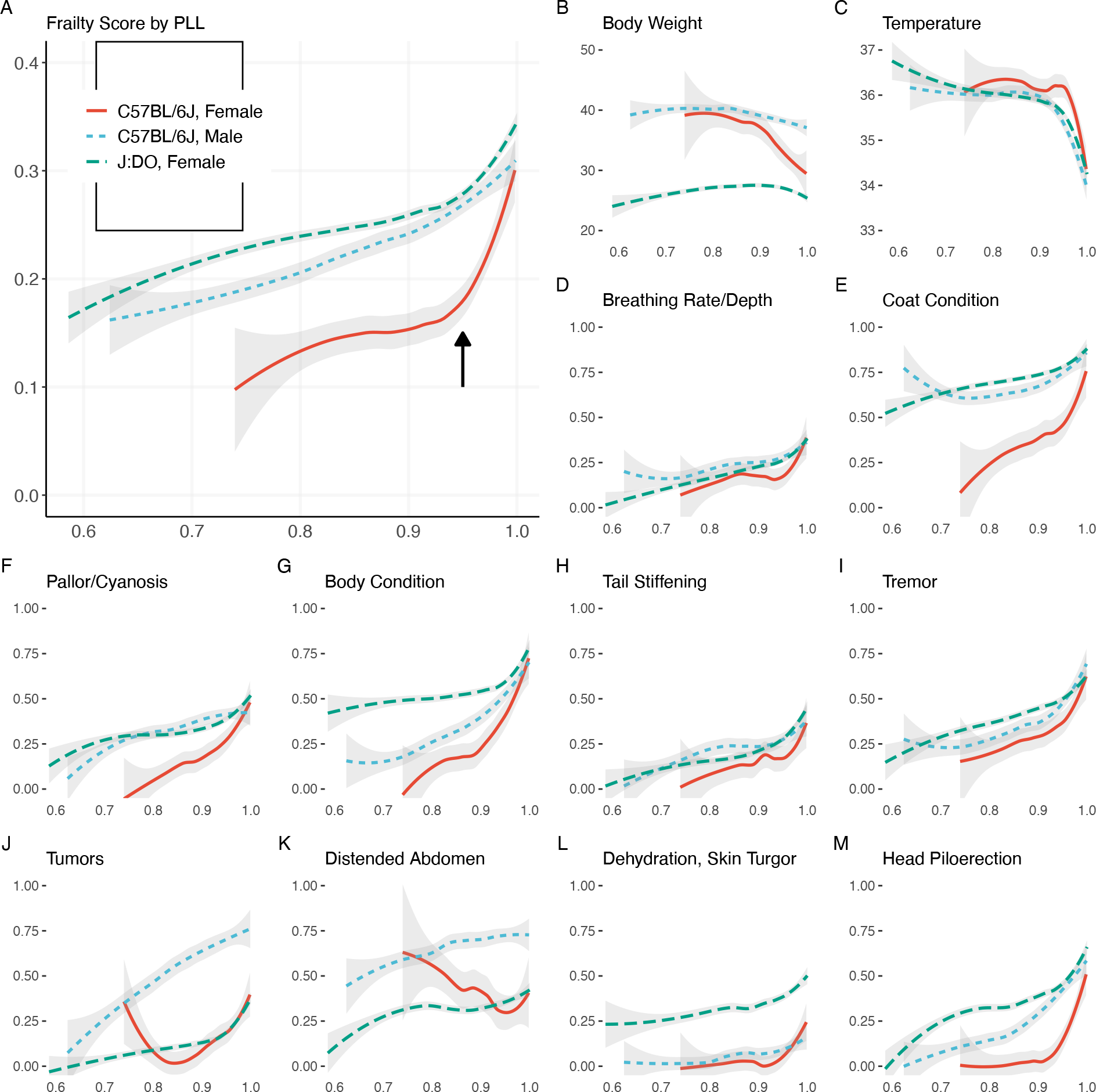
Longitudinal frailty assessments reveal inflection point near 0.95 PLL in DO females. X-axis: PLL range observed, 0.6 to 1.0. Y-axis: Locally weighted smoothed mean frailty. The 12 individual frailty items highlighted showed strongest correlation with life expectancy in the J:DO cohort (correlation data shown in Figure 1). The selected items illustrate a range of PLL responses; remaining items displayed in eFigure 1. J-shaped curves observed in mean body condition score and mean palpable mass score among female C57BL/6J mice were statistical artifacts. Re-evaluation with smaller smoothing span revealed patterns more consistent with generally increasing prevalence.

Plot of temperature trajectories on the PLL scale show a similar inflection point (**Figure 2d**). Examination of individual deficits on the PLL scale (**Figure 2e**, **Figure 2f**) reveals heterogeneity of lifetime trajectories across genetic background and sex. For example, clinical presentation of head piloerection among B6 females was unlikely until 0.9 PLL which is later than for other study strata.

Correlation coefficients for life expectancy with health deficits by genetic background (**Table 1**) identify candidate items for lifespan prediction in each genetic background. As in Schultz *et al*. [16], chronological age was a better predictor of life expectancy than individual health deficits among B6 mice. Yet among DO females, breathing rate/depth scores and palpable mass scores were more strongly associated with life expectancy than age. Among B6 mice, body condition score was more strongly associated with life expectancy than FgI score. The relationship between age and life expectancy was stronger and more precise among B6 mice (r [95% CI]: -0.457 [-0.507, -0.403] than DO mice (r [95% CI]: -0.209 [-0.236, -0.183]), likely due to lesser variance in lifespan in the inbred mice. Comparisons by diet among DO mice (**eTable 4**) and by sex among B6 mice (**eTable 5**) showed notable differences. For example, body condition, body weight, gait disorders, and tumors were more strongly associated with life expectancy among AL than CR40 female DOs, while FgI score and age were more strongly associated with life expectancy among CR40 than AL female DOs.

**Table 1:** Correlation coefficients (95% confidence intervals) and p-values for life expectancy with fragility items.

|  | J:DO | p | C57BL/6J | p |
| --- | --- | --- | --- | --- |
| Fgl Score | -0.27 (-0.29,-0.24) | <0.001 | -0.31 (-0.37,-0.25) | <0.001 |
| N Deficits | -0.24 (-0.27,-0.22) | <0.001 | -0.26 (-0.32,-0.19) | <0.001 |
| Breathing Rate/Depth | -0.23 (-0.25,-0.2) | <0.001 | -0.16 (-0.22,-0.09) | <0.001 |
| Age in weeks | -0.21 (-0.24,-0.18) | <0.001 | -0.46 (-0.51,-0.4) | <0.001 |
| Tumors | -0.21 (-0.23,-0.18) | <0.001 | -0.20 (-0.26,-0.14) | <0.001 |
| Temperature | 0.20 (0.17,0.23) | <0.001 | 0.24 (0.17,0.3) | <0.001 |
| Head Piloerection | -0.20 (-0.23,-0.17) | <0.001 | -0.32 (-0.38,-0.26) | <0.001 |
| Tail Stiffening | -0.19 (-0.22,-0.17) | <0.001 | -0.14 (-0.2,-0.08) | <0.001 |
| Tremor | -0.19 (-0.22,-0.17) | <0.001 | -0.28 (-0.34,-0.22) | <0.001 |
| Coat Condition | -0.17 (-0.2,-0.15) | <0.001 | -0.07 (-0.13,0) | 0.04 |
| Pallor/Cyanosis | -0.14 (-0.16,-0.11) | <0.001 | -0.19 (-0.25,-0.12) | <0.001 |
| Body Condition | -0.11 (-0.13,-0.08) | <0.001 | -0.35 (-0.4,-0.29) | <0.001 |
| Distended Abdomen | -0.11 (-0.14,-0.08) | <0.001 | -0.04 (-0.1,0.03) | 0.27 |
| Dehydration, Skin Turgor | -0.11 (-0.14,-0.09) | <0.001 | -0.13 (-0.2,-0.07) | <0.001 |
| Dermatitis | -0.10 (-0.13,-0.07) | <0.001 | -0.09 (-0.16,-0.03) | 0.01 |
| Urine | -0.10 (-0.13,-0.07) | <0.001 | -0.10 (-0.16,-0.03) | <0.001 |
| Thoracic Mass | -0.09 (-0.11,-0.06) | <0.001 | -0.03 (-0.09,0.04) | 0.43 |
| Gait Disorders | -0.08 (-0.11,-0.05) | <0.001 | -0.10 (-0.17,-0.04) | <0.001 |
| Body Weight | -0.07 (-0.1,-0.04) | <0.001 | 0.19 (0.13,0.26) | <0.001 |
| Paralysis | -0.07 (-0.1,-0.04) | <0.001 |  |  |
| Rectal Prolapse | -0.06 (-0.08,-0.03) | <0.001 | -0.11 (-0.17,-0.04) | <0.001 |
| Eye Discharge/Eyelid Inflammation | -0.05 (-0.08,-0.02) | <0.001 | -0.16 (-0.23,-0.1) | <0.001 |
| Kyphosis | -0.04 (-0.07,-0.01) | 0.01 | -0.07 (-0.13,0) | 0.04 |
| Hunched | -0.04 (-0.07,-0.01) | <0.001 | -0.22 (-0.28,-0.16) | <0.001 |
| Changes to Eye Globe | -0.04 (-0.07,-0.01) | <0.001 | -0.07 (-0.13,0) | 0.04 |
| Response to Analgesic | -0.04 (-0.07,-0.02) | <0.001 |  |  |
| Peri-retro-orbital Swelling | -0.03 (-0.06,0) | 0.02 | -0.06 (-0.12,0.01) | 0.1 |
| Activity | -0.03 (-0.06,0) | 0.03 | 0.05 (-0.01,0.12) | 0.1 |
| Malocclusions | -0.03 (-0.05,0) | 0.06 |  |  |
| Piloerection | -0.02 (-0.05,0.01) | 0.12 | 0.07 (0.01,0.14) | 0.03 |
| Diarrhea | -0.02 (-0.05,0) | 0.09 | 0.03 (-0.04,0.09) | 0.38 |
| Response to External Stimuli | -0.02 (-0.04,0.01) | 0.25 |  |  |
| Vestibular Disturbance | -0.00 (-0.03,0.03) | 0.98 | 0.06 (-0.01,0.13) | 0.07 |
| Nasal Discharge |  |  |  |  |
| Vaginal/Uterine Prolapse |  |  |  |  |

Deficits ordered by descending absolute correlation coefficient in J:DO cohort.

### Does the TxW criterion for imminent terminal decline generalize across experimental conditions, genetic backgrounds, and sex?

A lifespan study in 110 outbred CD1 mice found that a simple derived score, temperature x body weight (TxW), could identify mice in the last 2 to 3 weeks of life [15]. This was done by comparing the mean value of TxW in the most recent 2 to 3 weeks to a benchmark mean of values collected 10 to 13 weeks prior to current date. Greater than 10% reduction in TxW was taken to indicate imminent terminal decline. Because one week was roughly equivalent to 1% of lifespan in the study, preemptive euthanasia guided by TxW would potentially have resulted in 2% underestimation in total survival time. The simplicity of the metric, the outbred status of the mice, and the 100% precision of this criterion all indicated promise for the TxW score. However, the criteria were derived post hoc, without internal validation to control the risk of overfitting the predictive endpoint classifier. In a later study, TxW did not appear to improve sensitivity or specificity of LLD detection in AKR/J, C57BL/6J, or BALB/cByJ mice [18]. The investigators recognized these issues and encouraged others to evaluate the TxW metric in their own studies. Prediction models generally perform less well in external validation than in development [22].

As in Ray *et al*. [15], timeseries data for the product of weight and temperature within mice were separated into moving average intervals to identify sustained reduction of weight and temperature. Ten percent reduction in mean TxW from the first 4 weeks to the last 2-to 3-weeks in a 3-month interval indicated terminal decline. We estimated predictive performance of TxW within genetic backgrounds, diets, and sex. We considered both the positive predictive value (PPV is the probability that, following a positive prediction, the sample is truly positive) and the negative predictive value (NPV is the probability that, following a negative prediction, the sample is truly negative) of the TxW criterion. The results (**eTable 6**) reveal high negative predictive value (NPV) but low positive predictive (PPV) in our data for all strata except DO mice assigned IF treatment. Low PPV could lead to false positive predictions of LLD, premature euthanasia, and a consequent downward bias in estimates of lifespan.

### Can weekly clinical assessments in aged mice improve prediction of imminent decline?

We next aimed to produce a binary classifier capable of predicting imminent terminal decline in DO mice, capitalizing on the over 5,000 quantitative assays in the DO mice (bodyweight [grams], temperature [Celsius]) supplemented by weekly assessments using our FgI. We defined imminent decline as within the last 5% of lifespan, or 95% proportion of life lived (95PLL). Detailed methods including data pre-processing steps, train/test set splitting, v-fold cross-validation, model stacking, and feature performance are provided in Methods. In an effort towards interpretable ML, we inspected stacking coefficients used to weight predictions from candidate models in an ensemble and SHapley Additive explanation (SHAP) visualization plots of the highest weighted candidate model to quantify feature/predictor importance in test data. Stacking coefficient ranks (**eFigure 2a**) and variable importance plots (**eFigure 2b, eFigure 2d**) indicated age and dietary assignment were critical to the ensemble’s 95PLL prediction in these data. Without pre-specifying non-linearity or interaction effects, dependence plots revealed nonlinear associations between age in weeks and 95PLL (**eFigure 2c**) and nonlinear associations between FgI score and 95PLL (**eFigure 2e**), and interaction plots identified modifying effects between FgI items and age on 95PLL risk (**eFigure 2f**).

We applied this ML model to the held-out test data to predict 95PLL based on the features extracted from the new data. The model achieved an accuracy of 0.79, meaning it correctly classified four of every five cases. The positive predictive value (PPV) and negative predictive value (NPV) of the model indicated that ∼ 60% of the positive predictions and ∼ 80% of the negative predictions were true. The area under the precision-recall curve (PR-AUC) and the area under the receiver operating characteristic curve (AUC) of the model were 0.61 and 0.80, respectively. The AUC is a measure of how well a model can distinguish between different classes, i.e., whether mice have reached 95PLL or not. The closer the AUC is to 1, the better the model. These metrics suggest that the model performed well in predicting 95PLL status, but there is still room for improvement. Binary classification models such as this one determine Yes/No outcome prediction based on the predicted probability of an event. Typically, the threshold is set at 50%, but the cutoff probability can be set higher or lower, depending on the context. We explored various predictive probability thresholds and found that the ML-FgI ensemble model correctly flagged approximately one-third of DO mice at 95PLL with minimal risk of a false positive when predicted probabilities of 95PLL status were at or above 0.6.

To facilitate comparison across different lifespan predictors, we adapted the TxW criteria to predict 95PLL. We fit a generalized linear mixed-effects model within strain strata to estimate risk of 95PLL from TxW status (pass/fail) as a fixed effect and subject ID as a random effect in the training data. The predicted probability of 95PLL was estimated using the fixed effects term for TxW in the testing data (*(β*_TxW_ AUC = 0.649 in C57BL/6J; *(β*_TxW_ AUC = 0.573 in J:DO). We also fit a generalized linear mixed-effects model with TxW that incorporated the FgI score as a continuous variable with range 0 to 1. The TxW+FgI model represents an intermediate level of complexity between TxW and the ML-FgI ensemble model. One potential advantage of the TxW+FgI model is that it uses only the FgI and does not depend on a specified set of items. Within strain strata, a formula estimating risk of 95PLL from TxW status (pass/fail) and FgI score (mean of 30 ordinal items) as fixed effects and subject ID as a random effect was fit in training data. To evaluate predictive performance of the TxW+FgI models generated, predicted probability of 95PLL was estimated using fixed effects terms from the fitted model in testing data (*(β*_TxW_+*(β* _FgI_ AUC = 0.765 in C57BL/6J; *(β*_TxW_+*(β* _FgI_ AUC = 0.658 in J:DO).

The ranked performance of the DO-specific models based on AUC (ML-FgI *>*TxW+FgI *>*TxW; **Figure 3**) followed increasing computational complexity of the models; it showed that inputting cross-sectional data from weekly frailty phenotyping substantially improved performance relative to TxW (TxW+FgI *>*TxW); and it revealed that inputting the temporal dynamics of densely sampled longitudinal frailty phenotype as 30-day differences could further improve performance relative to cross-sectional phenotypic descriptors (ML-FgI *>*TxW+FgI).

**Figure 3:**
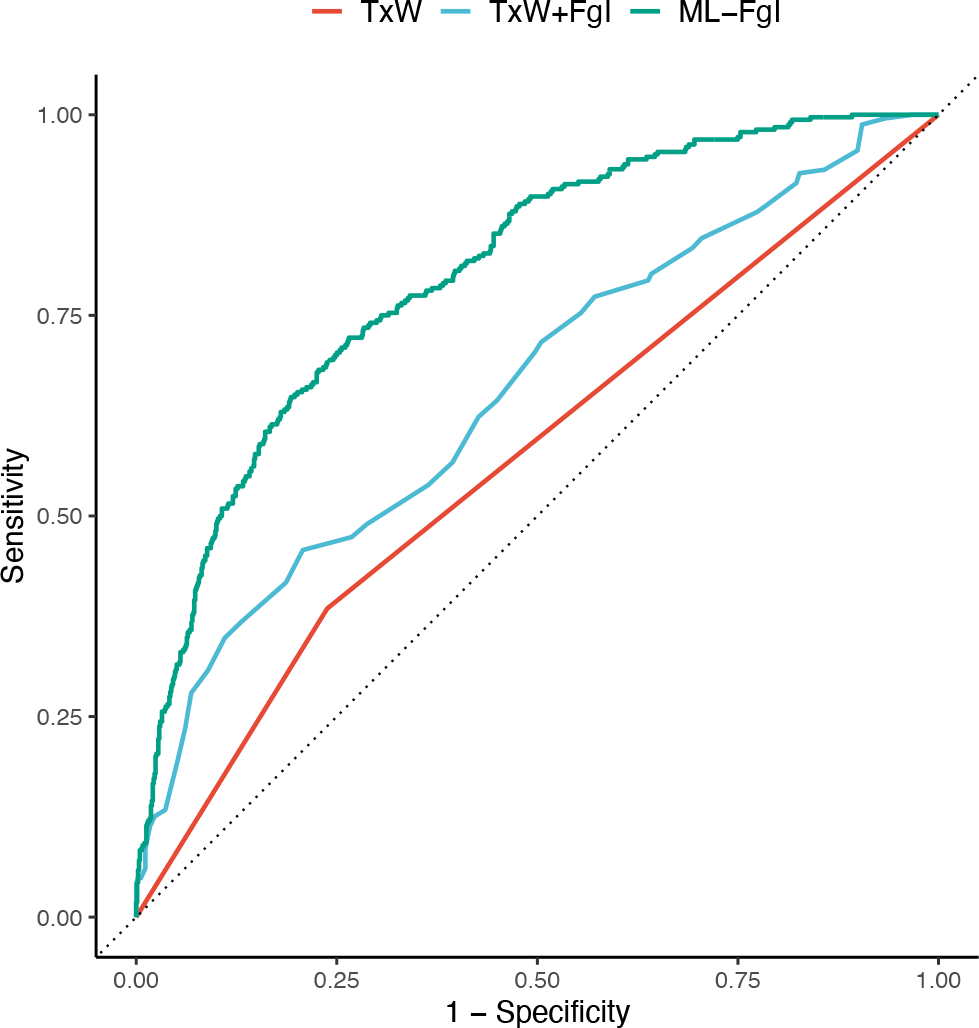
Longitudinal frailty assessments improve predictive endpoint performance in DO females. A) J:DO = diversity outbred. ROC=receiver operating curve. The TxW predictive endpoint uses a simple algebraic comparison of weekly temperature and body weight history over previous 3 months. The TxW+FgI predictive endpoint uses a simple algebraic expression of TxW status (pass/fail) and most recent summary FgI score. The ML-FgI ensemble predictive endpoint uses a complex ML algorithm to identify patterns in 30 densely sampled binarized longitudinal frailty phenotypes (most recent and 30-day subsequent differences), body weight (30-day subsequent difference), and temperature (30-day subsequent difference), sex, and diet. See methods for model development methods. Performance was calculated on testing datasets specific to genetic background. Data were randomly allocated to training and testing datasets by mouse rather than by assessment. A probability threshold of 0.6 minimized the false positive rate to 5% while still identifying one-third of the true positives in the test data.

### Frailty-based healthspan - an alternative target for preclinical aging research

When considering the efficacy of aging interventions, the most desirable outcome is to extend the length of time lived in good health. We considered using the FgI to derive a measure of healthspan as an alternative outcome for aging studies. We defined the healthspan endpoint as the time of occurrence of four or more severe health deficits as assessed by our FgI. A similar definition of healthspan was previously proposed [23]. We chose this definition because almost all animals in the study population experienced it at some point during the follow-up period. This ensured that the censoring rate was minimal and that the survival curves reflected the true distribution of the event times. Healthspan was thus defined as length of life lived with fewer than four severe health deficits. Most mice met this threshold, although proportionally more aged DO mice met this threshold prior to exit (99%) than aged B6 mice (89%). Median [Q1, Q3] healthspan among B6 mice was 24.9 [24.2, 26.3] months (p sex effect=4.7e-5) and for DO mice was 31.6 [31.6, 32.09] months (p diet effect=0.076). Maximum healthspan, estimated as the 90th percentile of healthspans within a genetic background was 27.9 months among B6 mice (p sex effect = 0.10) and 38.5 months among DO mice (p diet effect = 0.39).

Survival distributions for healthspan and lifespan suggest that, as in humans [24], lifespan extension factors and healthspan extension factors are distinct (**Figure 4**). Differences in lifespan observed by diet among DO females were less evident in the healthspan metric, particularly at earlier ages, and although sex differences in lifespan were not observed in the C57BL/6J mice, sex effects on healthspan were strong, favoring longer healthspan among the female C57BL/6J mice. Groupwise comparisons of this healthspan measure can mirror lifespan extension intervention assessment: Two-thirds of the DO mice with maximum estimated healthspan came from the CR20 and CR40 diet groups, suggesting these diets may delay age-related decline by increasing maximum healthspan relative to IF or AL diets. Comparison of sex effects on healthspan among aged C57BL/6J mice suggested potential healthspan extension in females compared to males (quantile test p = 0.01).

**Figure 4:**
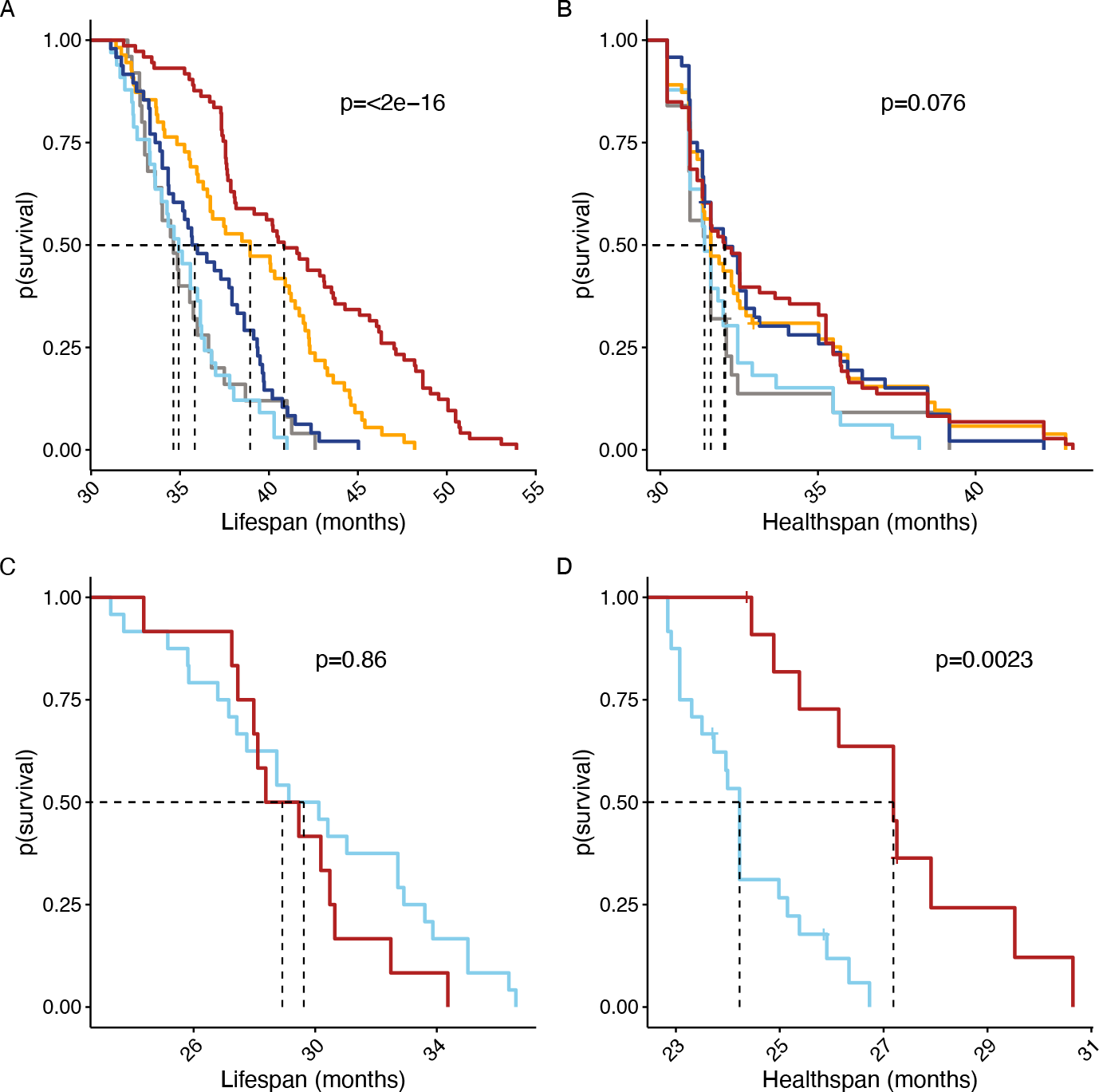
Lifespan and healthspan survival curves. Lifespan and healthspan by randomization group among DO female mice (A and B, color legend as in Figure 1) or by sex among C57BL/6J mice (C and D, color legend as in Figure 2. Log-rank p-values compare lifespan (A and C) and healthspan (B and D) across groups within strain cohort. Healthspan was defined as length of life lived with fewer than four severe health deficits. Vertical lines indicate median survival.

## Discussion

Aged mouse models are often used to characterize normative aging (as in the Study of Longitudinal Aging in Mice [14], to evaluate purported lifespan interventions (as in the Intervention Testing Program [25]), and to unravel agespecific disease mechanisms (as part of the National Institute on Aging’s Nathan Shock Center initiative [26–30]. These studies rely on natural lifespan as their primary outcome. The ability to predict lifespan could reduce the time and costs associated with these studies as well as the need for late-life veterinary intervention. Swindell *et al*. [3] presented an early attempt to achieve lifespan prediction using ML techniques applied to serum biomarkers and other traits. They concluded that “… such experiments [full-length survivorship] will never be replaced by predictive models. Well-developed models, however, can generate preliminary data years in advance, and the output of such models, as an integration of multiple variables that may predict life span individually, can provide a surrogate target for aging research that is easier to evaluate than lifespan.” [3]

To address the unmet need for validated predictive endpoints in longevity and aging research, we established a large, aged, outbred cohort of mice and densely sampled bodyweight, temperature, and frailty over multiple years. This resource was later enhanced by the addition of an aged, inbred mouse cohort. We aimed to evaluate the utility and limitations of easily assessed biomarkers for predicting lifespan, healthspan, and terminal decline in longitudinal aging studies in mice. We used observable clinical markers to construct a Fragility Index (FgI) based on the human frailty index [31]. We chose this approach because it was less invasive than quantitative assessments of health (e.g., serum biomarkers), it was easy to implement, and could be repeated at high frequency (weekly).

We first examined an existing criterion for predicting imminent terminal decline in longitudinal aging studies with mice based on temperature and body weight (TxW). We found that, while TxW performed substantially better than chance (AUC ∼= 0.6), it did not achieve the high level of predictive accuracy that would be needed to support preventive euthanasia in this multi-factorial longitudinal cohort study.

We then incorporated data from the FgI that had been conducted weekly from midlife (2 or 2.5 years) until end of life. We adopted ML methods to analyze these data. The best performance was achieved using an ensemble model that stacked candidate models instead of choosing the single best-performing model. Our ensemble algorithm represented an improvement over previous predictive criteria but fell short of the level of reliability needed to accelerate lifespan studies. The ensemble model is complex to implement and, as with the AFRAID algorithm [16], relies on the evaluation of a specific set of items in a frailty index. One of the appealing properties of FIs is that there is some flexibility in the choice of items to score. To obtain a more portable predictor, allowing for some variation of the items that make up the FI, we used only the summary FgI score in our TxW+FgI predictor, which demonstrated an intermediate level of performance. Taken together, our ensemble algorithm performance, and previous research [19] suggest that a generalizable predictor of lifespan derived from FIs may be at an impasse.

While frailty-based prediction falls short as a surrogate for lifespan, it did demonstrate significant predictive power and as such must contain information that could be used to inform the conclusion of aging experiments. This led us to examine the utility of FIs as a measure of healthspan. We constructed a simple definition of healthspan based on the number of items scored as severe [23]. We found that the effects of factors such as sex (in the C57BL/6 mice) and diet (in the DO mice) that are known to affect lifespan do not necessarily have the same effect on healthspan. The lack of correlation between healthspan and lifespan has been previously reported [32]. For example, the combined sex effects of abbreviated lifespan and extended healthspan observed among aged female C57BL/6J mice relative to aged male C57BL/6J mice suggests that aged B6 females presented a compressed morbidity period [33] relative to males in this study. Refocusing the field’s priorities from lifespan to healthspan as primary outcome may have benefits such as increased clinical relevance.

There is no consensus on how to define and measure healthspan in animals. In this study, we computed healthspan according to a frailty-threshold definition. Frailty-adjusted Mouse Years (FAMY) are another recently developed definition of healthspan with the potential for high clinical translatability as a preclinical analog to QALYs [34]. Any definition will have advantages and limitations for assessing healthspan. FAMY computations assume a linear relationship between time and utility, which may not reflect the true valuation of time in different stages of life and health. The frailty-threshold approach is a simpler definition of healthspan, but this simplicity comes at the cost of coarser data: healthspan is linked to frailty collection dates, which are not usually collected as densely as in the Fragility Longitudinal Study.

We aimed to understand the aging process in mice across different genetic backgrounds, sexes, and experimental conditions. High dimensional datasets like the one generated here allow us to arrive at fine-grained understanding of the aging process. While the collection of large numbers of phenotypes is increasingly common, this study is distinctive for the temporal density of those phenotypes (weekly), for using a highly genetically diverse cohort of outbred mice, for including both males and females, and for including a control cohort of inbred mice. As in humans [35], we find that single summary measures of aging such as TxW do not predict individual outcomes (such as 95PLL) in a highly heterogenous populations. These findings have important implications for aging research outcome priorities. Lifespan in mice is largely determined by neoplastic disease risk and using lifespan as the key metric for geroprotective intervention may limit anti-aging interventions developed in mice to anti-cancer effects [36]. Aging research addressing factors that impact quality of life and healthspan [37] broadens therapeutic targets for anti-aging treatment in murine models and in humans. In the future, metabolomics, proteomics, and other quantitative biomarkers might provide insight into how frailty manifests as a first step towards developing interventions targeting healthspan by decelerating frailty accumulation. Such developments would have high translational value regardless of their impact of lifespan.

Since our study was initiated, there have been significant advances in non-invasive methods for frailty assessment in mice and humans. For example, machine-vision-based FI for mice using video cameras and ML to automate frailty index scoring from morphometric features, behavioral assessment, and gait analysis [38, 39]. These methods could enable high-throughput frailty phenotyping in large colonies of aged mice without overwhelming human labor resources. Similarly, wearable technology and sensor-based mobility assessments have been used to measure frailty in humans [40, 41]. These co-occurring innovations may result in improved equivalence across human and mouse models of health deficit accumulation in aging. They may also increase experimental power and reduce the number of experimental animals required for statistical significance, in keeping with the 3Rs principle of Reduction. However, these methods may also require strain-specific calibration, as machine vision indices appear to vary across different mouse strains [42]. The current study could be reimplemented employing these newer technologies for high throughput frailty phenotyping, which geriatric experts have highlighted as an important advancement for the field [41].

The predictive models developed and evaluated in this study may serve as a tool to guide timing and intensity of clinical monitoring but cannot adequately predict lifespan for the purpose of prescribing preventive euthanasia. A generalized predictive endpoint at the level of the individual for aging and longevity research in mice with high sensitivity and high specificity to replace lifespan as a primary outcome remains an unmet need. An alternative to prediction could be to merge findings from lifespan and healthspan to generate more comprehensive assessments of anti-aging therapeutics. The field may benefit from an increased focus on healthspan to better characterize the heterogeneity of aging trajectories and increase the clinical relevance of aging studies in animal models.

## Materials and Methods

### Mice

A small proportion of mice (6.6%) with missing natural lifespan data were excluded from the study. This could be due to failure to recover from anesthesia conducted as part of another study protocol or due to death type missing or discarded, likely reflecting the epidemiology of fighting among mice living together in long-term groups under standard laboratory housing conditions [43]. All mice were maintained on a standard 6% fat/kcal chow diet (Lab Diets 5KOG), with a 12:12 light dark cycle and room temperature 72F.

Research staff regularly evaluated mice for pre-specified clinical signs in addition to the FgI assays: palpable hypothermia, responsiveness to stimuli, ability to eat or drink, dermatitis, tumors, abdominal distention, mobility, eye conditions (e.g., corneal ulcers), malocclusion, trauma, and wounds of aggression. If mice met the criteria for observation in any of these categories, veterinary staff were contacted. If the clinical team determined a mouse to be palpably hypothermic and unresponsive, unable to eat or drink, and/or met protocol criteria for severe dermatitis, tumors, and/or fight wounds, preemptive euthanasia was performed to prevent suffering; otherwise, the veterinary staff provided treatment. The goal of the study was to predict mice that would meet these criteria before developing a moribund condition.

All procedures used in this study received prior approval from the Jackson Laboratory’s Animal Care and Use Committee under protocol #06005.

### Phenotypic Assays

The Fragility index (FgI), developed to quantify the accumulation of detrimental health outcomes with age [31]. Several FI have been proposed; these are typically comprised of non-invasive and easily observable phenotypes associated with the aging process. Previous work using C57BL/6 mice demonstrated that an FI can quantify mortality risk in mice similar to mortality risk assessed by deficit accumulation in humans [44]. This and other studies [45] established merit for using FI in preclinical models of aging and in prediction of LE. Modified clinical frailty assessments were derived from existing FI [13] with additional deficits selected by clinical staff based on their years of experience in the evaluation and treatment of mice in aging studies. Modified items in the FgI concerned features of integument (new: pallor and/or cyanosis; not assessed: alopecia, loss of whiskers, loss of fur color), physical/musculoskeletal health (new: palpable thoracic mass; split: kyphosis into kyphosis and hunched posture; not assessed: forelimb grip strength), vestibulocochlear health (not assessed: hearing loss), ocular/nasal health (new: peri-retro-orbital swelling and response to analgesic; collapsed: corneal opacity, cataracts and micropthalmia into changes to eye globe; not assessed: vision loss and menace reflex), digestive/urogenital health (new: urinary expression and dehydration), discomfort (split: piloerection into head piloerection and other; not assessed: mouse grimace scale), and behavior (new: activity levels, responsiveness to external stimuli, and paralysis).

The 32-deficit assessment consisted of two quantitative assays (bodyweight [grams] and rectal temperature [Celsius]) and 30 ordinal assays (**eTable 1**). Most ordinal assays were rated on a 3-level scale as absent, mild, or severe. Exceptions were rated on a 2-level scale as absent or severe. Further granularity in the assessments was deemed to be overly time-consuming and potentially error prone [46]. Only assays that could be carried out in typical preclinical laboratory research settings were considered for inclusion. All mice enrolled in the study and still living were evaluated weekly. Most assessments were completed by one of two technical experts who conducted 66% (n=4,200 indices) and 30% (n=1,855 indices), respectively. The remaining 4% of assessments were completed by two other technicians. The order of assays was consistent: bodyweight recording, followed by one hour of habituation, followed by the ordinal observational assays and then body temperature measurement. Two deficits (head piloerection and thoracic mass) were both missing among 5% of assays; these two deficits were added to the index in the third month of the study based on observations made by study staff. Of the remaining 30 deficits, data were either completely observed or *>*99.9% complete. One mouse that achieved very late life (4.5 years) was switched to a monthly phenotyping at approximately 4.4 years of age. In December 2021, two mice remained in the study, and research staff halted assessments for the remainder of life (1 and 2 months).

## Data Processing

### Data Cleaning

Initial data quality control included identifying and resolving equipment mis-calibration, mislabeled animals, and technically impossible values (e.g., 0 degrees Celsius temperature for a living animal). For ordinal variables, out of range values were identified and resolved based on technician notes. Non-meaningful zeros were resolved as missing data. FgI data collection relied on paper-based recording and manual data entry into a laboratory information management system that allowed for quality control but was not real-time. We note that computerized data collection with real-time quality control can improve assessment quality and should be considered in future studies [47].

### Computing TxW criteria

As in Ray *et al*. [15], timeseries data for the product of weight and temperature within mice were separated into moving average intervals to identify sustained reduction of weight and temperature. Ten percent reduction in mean TxW from the first 4 weeks to the last 2-to 3-weeks in a 3-month interval indicated terminal decline.

### Computing FgI score

Most deficits were scored on a two-[0,1] or three-[0,0.5,1] level ordinal scale where 0 indicated the absence of the health status deficit; 0.5 indicated mild deficit; and 1 indicated severe deficit. Dermatitis was an exception with four levels but was simplified to three levels during data processing. Thus, all deficits in the index were established as ordinal values prior to computing a mean, yielding an FgI score between 0 and 1 for each FgI assessment.

### Statistical Methods

We carried out separate parallel analyses for B6 and DO data. To determine the cumulative incidence of each deficit, a maximum deficit score was computed for each mouse. The proportion of mice that displayed a maximum possible deficit score was computed by deficit across each aged cohort. Cumulative incidence was used to reveal ceiling and floor effects in the health deficit scale. Descriptive summary statistics for FgI score were computed to provide benchmarks that can be used by the pre-clinical aging research community to guide study design and planning. Lifespan in the Longitudinal Fragility Study was compared by t-tests and one-way ANOVA. A test for homogeneity of variance across genetic backgrounds was computed by comparing deviation scores to group medians. To compare distributions in maximum lifespan, estimated as the upper 90th percentile weeks of the lifespan distribution, quantile tests of proportions were estimated. Healthspan was defined as length of life lived with fewer than four severe health deficits; this is a post-hoc threshold selected for near census occurrence in the two cohorts. Healthspan data were compared as in the mortality data, via score tests and quantile tests of proportions. To identify inflection points in health status over residual lifespan, FgI score and proportion of life lived (PLL = current age divided by lifespan) were visually displayed as smoothed local polynomial regression fitting.

To validate the TxW criteria [15], we computed the positive and negative predictive values (PPV and NPV respectively) within genetic backgrounds, diets, and sex. PPV is defined as the proportion of assessments that had reached the last 5% of lifespan (*>*95% PLL) at the time of positive prediction. NPV is the proportion of assessments at *<*=95% PLL at the time of a negative prediction. In the current study, high PPV would indicate that early euthanasia would result in underestimation of natural lifespan by less than 5%; high NPV would indicate that mice that are not euthanized early have not yet reached the final 5% of lifespan. Due to the differences between the FI employed by Schultz *et al*.

[16] and the FgI used in the current study, we could not pursue external validation testing of the AFRAID clock in these data.

Results from this study are based on exploratory analyses of observational data and should be considered preliminary; p values less than 0.05 were considered significant without adjustment for multiple comparisons. All estimates should be interpreted as conditional on survival to study enrollment. Analyses were conducted using the R Statistical language (version 4.2.3; [48]).

### Machine Learning

We aimed to produce a binary classifier capable of predicting imminent terminal decline from baseline characteristics and dense longitudinal data in DO mice. The following is a detailed description of the data pipeline, including data pre-processing steps, train/test set splitting, v-fold cross-validation, model stacking, and comparative performance.

We operationalized terminal decline as within the last 5% of lifespan, or 95% proportion of life lived (95PLL). The objective function optimized was the area under the receiver operating curve (AUC-ROC). Feature sets included known lifespan predictors: age, diet [49], temperature [50], body weight [8], and frailty [44]. We developed predictive endpoint classifiers based on four different feature sets: model 0 included age at assay, and diet; model 1 included model 0 + temperature and body weight, raw and pre-post percent difference for first assessment in 30 days prior to the current test vs. current assessment; model 2 included model 1 + 30 ordinal FgI deficit items; model 3 included model 2 + 30-day pre-post difference (computed as in temperature and body weight). DO mice with *<*30 days of data were excluded (n=29). Of the remaining DO assessments (n=4995), 276 assessments (5.5%) contained missing data which were excluded. ML classifiers perform better on balanced datasets. Where data are unbalanced, outcome classes can be up-sampled or down-sampled to make the occurrence of outcome classes equal [51]. In the current dataset, where events outnumbered non-events, down-sampling would remove rows of non-events and up-sampling would replicate event rows. To prevent data destruction, event rows were sampled up to have the same frequency as the most occurring level.

We applied data pre-processing steps to predictors in preparation for modeling as follows. Rolling window 30-day percent change in body weight and temperature were derived comparing current measurement (last in window) to the earliest available measurement in the previous 30 days (first in window). Rolling over a date index, versus n assessments, is useful to ensure equidistant data windows regardless of data missingness. Where current ordinal data from the 30 frailty deficits were entered into a model, these data were converted into one or more numeric binary model terms for the levels of the original data. Variables that contained only a single value (i.e., were of constant variance), were automatically removed. The 30 binary-encoded frailty deficits were translated from raw deficit scores having values ∈ {0, 1} or ∈ {0, .5, 1} (see above) to integer scales, either ∈ {1, 3} or ∈ {1, 2, 3}, ensuring absent and severe deficit corresponds to an equal numerical scored distance for binary and ternary deficits then converted into principal components of which 5 were retained. We extracted 30-day frailty item deltas, also summarized into five statistically independent principal components each. Data were normalized to have a mean of zero and a standard deviation of one.

Train/test set splitting and v-fold cross-validation are key aspects of model training to avoid overfitting, which compromises prediction performance in external data. In initial train/test data partitioning, we withheld data attributed to 25% of the unique animal identifiers to use later in evaluation of candidate models identified through model performance metrics in the training data stratified by diet. The training dataset was split into a series of 5 train-validate sets to improve the stability of model performance estimates in hyperparameter tuning. Candidate models were specified (max grid size per model = 25 hyperparameter combinations) and tuned across grouped resamples via a racing approach [52]. These procedures were repeated for each of four ML classifier model types (random forest [RF], support vector machines [SVM], regularized logistic regression [RLR], and eXtreme Gradient Boosting [XGB]), forming a large workflow set of four model specification by four preprocessors by many model configurations (n configurations determined by racing approach [above]).

Stacking candidate models instead of discarding all but the single best-performing model can boost model performance. A single ensemble model was derived from the workflow set in three steps. First, assessment set predictions across the v fold grouped splits were pooled into a single dataset. Second, a regularized linear model on the objective function computed coefficients for each entry in the workflow set/candidate ensemble member across the pooled dataset. Third, models with non-zero coefficients were retained to form a single ensemble model and trained on all training data from the initial grouped split. These fits plus the weighting coefficients determine the final ensemble model which was subsequently run on held out test data to estimate expected binary classification performance on new FgI assessments (i.e., performance evaluation). Performance metrics were computed across a series of alternate probability thresholds (0.5 to 0.9 by 0.1). Although they can be more computationally expensive and difficult to interpret than individual models, ensemble methods are often more robust to small changes in the training data or hyperparameter settings in any single predictive model. To aid in interpretability, SHAP (Shapley Additive exPlanations) values for the supervised ML model with the highest ensemble weight were computed [53]. ML interpretability can be used both as a hypothesis generator by identifying models not specified a priori and as a tool to guide future iterations of feature sets for predictive endpoints.

Comparative performance for the three predictive endpoints (TxW, TxW+FgI, ensemble) was assessed via ROC curve analysis in the DO test data. The estimated probability of an observation belonging to the 95PLL event class given its values on included features is referred to as the posterior class probability. We calculated the posterior probability of belonging to the 95PLL event class for each of the observations in the test set for the three predictive endpoints. When a predictor is categorical, the ROC curve has one less than the number of potential thresholds. A binary predictor as in TxW has only one threshold. To enable comparison of the binary TxW criterion against the two other predictive endpoint types via ROC, we modified TxW as follows: within strain strata, a formula estimating risk of 95PLL from TxW status alone as a fixed effect and mouse identifier as a random effect was fit in training data. To evaluate predictive performance of the modified TxW, predicted probability of 95PLL was estimated using a fixed effects term from the fitted model in testing data. Training and test sets were smaller for TxW and TxW+FgI than for ML-FgI because the TxW metric requires a longer look-back window. Sensitivity analyses for ROC curves comparing ML-FgI in the subset of testing data with complete TxW showed no difference in rank performance.

## Data Availability Statement

All analyses were performed using the R statistical programming language [48]. Code used to generate tables, figures, and reported results can be found on figshare (DOI: 10.6084/m9.figshare.25125587).

## Acknowledgments

We thank the JAX Nathan Shock Center Animal and Phenotyping Core team for their assistance with animal husbandry, data collection, and data curation. We thank the staff at Rockstep Inc., for their support with customization of our LIMS system, and Brenda Kick, The Jackson Laboratory, for helpful discussions and suggestions during study conception and design.

## Funding

This work was supported by the National Institutes of Health (Nathan Shock Centers of Excellence in the Basic Biology of Aging program, grant number AG38070 to R.K. and G.C.) and Calico Life Sciences LLC (Dietary Intervention of Aging in Genetically Diverse Animals, grant number CALICO-GAC-06 to G.C.).

## Disclosures

ADF is an employee of Calico Life Sciences LLC.

