## Supplemental Materials for "Longitudinal Fragility Phenotyping Predicts Lifespan and Age-Associated Morbidity in C57BL/6 and Diversity Outbred Mice"

**eFigure 1:** Locally weighted smoothed mean frailty by index item. X-axis: PLL range observed, 0.6 to 1.0. Y-axis: Locally weighted smoothed mean frailty. The 20 individual frailty items highlighted showed weak correlation with life expectancy in the DO cohort; remaining items and color legend displayed in Figure 2.

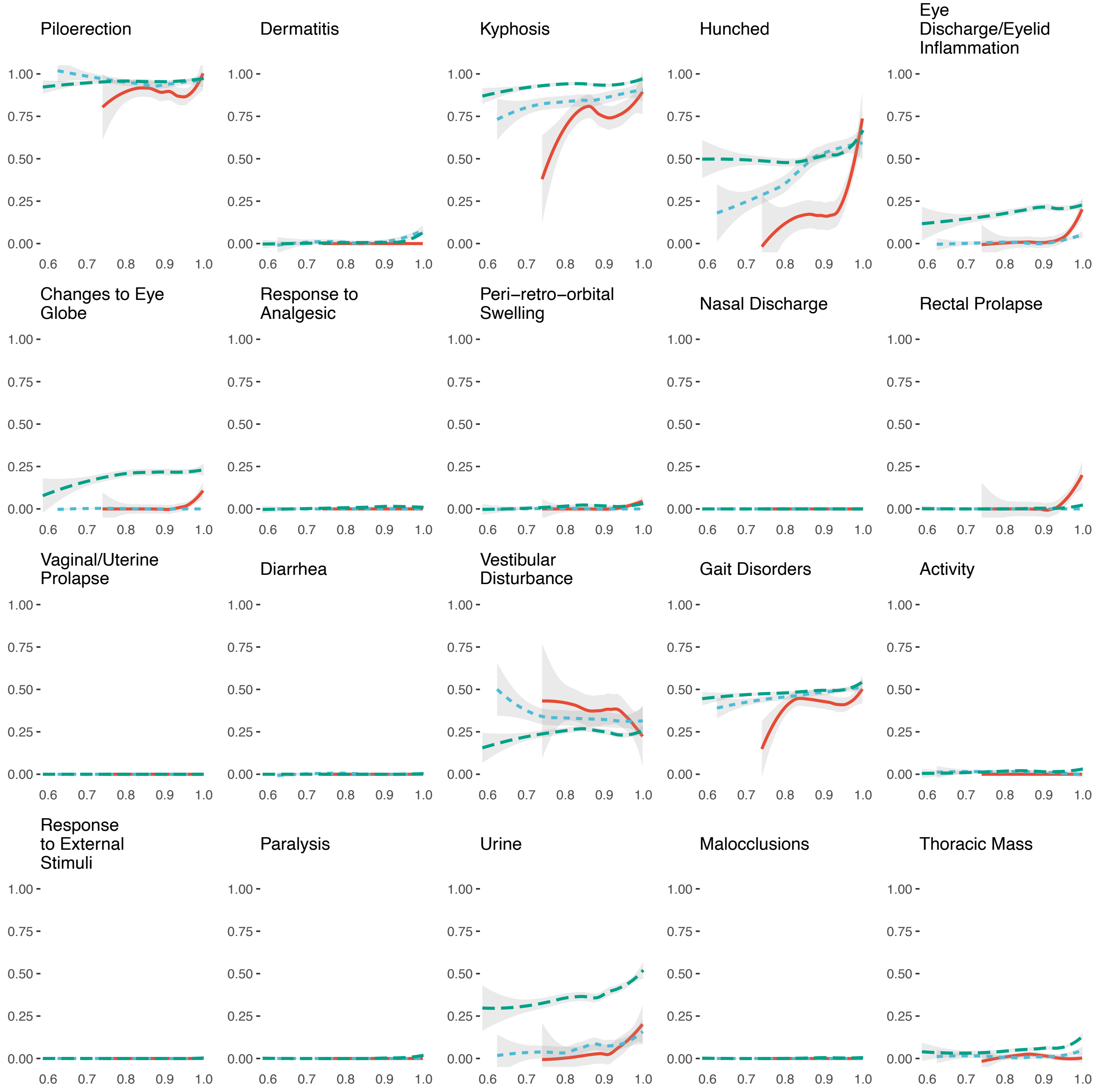

**eFigure 2:** Feature importance. A) Stacking coefficients are used to weight the predictions from each model in the ensemble. A parsimonious model with only age and dietary assignment was weighted highly. B) Variable importance*. Variables listed near the top of the figure are most important for 95PLL prediction in this model. C) Dependence plot*. Machine learning algorithm identified monotonic nonlinear association between age in weeks and 95PLL without pre-specifying non-linearity. D) Variable importance** E) Dependence plot**; Identified nonlinear relationship with frailty score where scores 2SD below mean are more informative than scores ˜-1.5SD below mean, and increasing scores are positively associated with importance thereafter. High frailty score (>=1.5SD) from mean is most informative in 95PLL prediction). F) SHAP interaction values for age and recent frailty change**. Visualizations generated with R package SHAPforxgboost Liu *et al.* [54]. *determined via SHAP values computed for XGB algorithm with the highest ensemble weight (rank 1; eFig 2A) **determined via SHAP values computed for XGB algorithm with a frailty feature with the highest ensemble weight rank (rank 3; eFigure 2A)

A

| Rank | Model | PP | Weight |
|---|---|---|---|
| 1 | XGB | m0 | 5.400 |
| 2 | RF | m3 | 1.431 |
| 3 | SVM | m1 | 0.915 |
| 4 | XGB | m3 | 0.732 |
| 5 | SVM | m3 | 0.729 |
| 6 | SVM | m1 | 0.340 |
| 7 | RF | m0 | 0.310 |
| 8 | SVM | m1 | 0.261 |
| 9 | SVM | m3 | 0.189 |
| 10 | SVM | m0 | 0.114 |

B

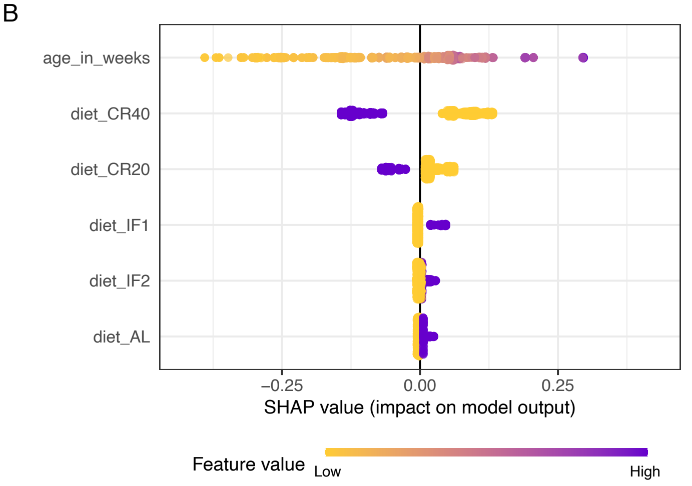


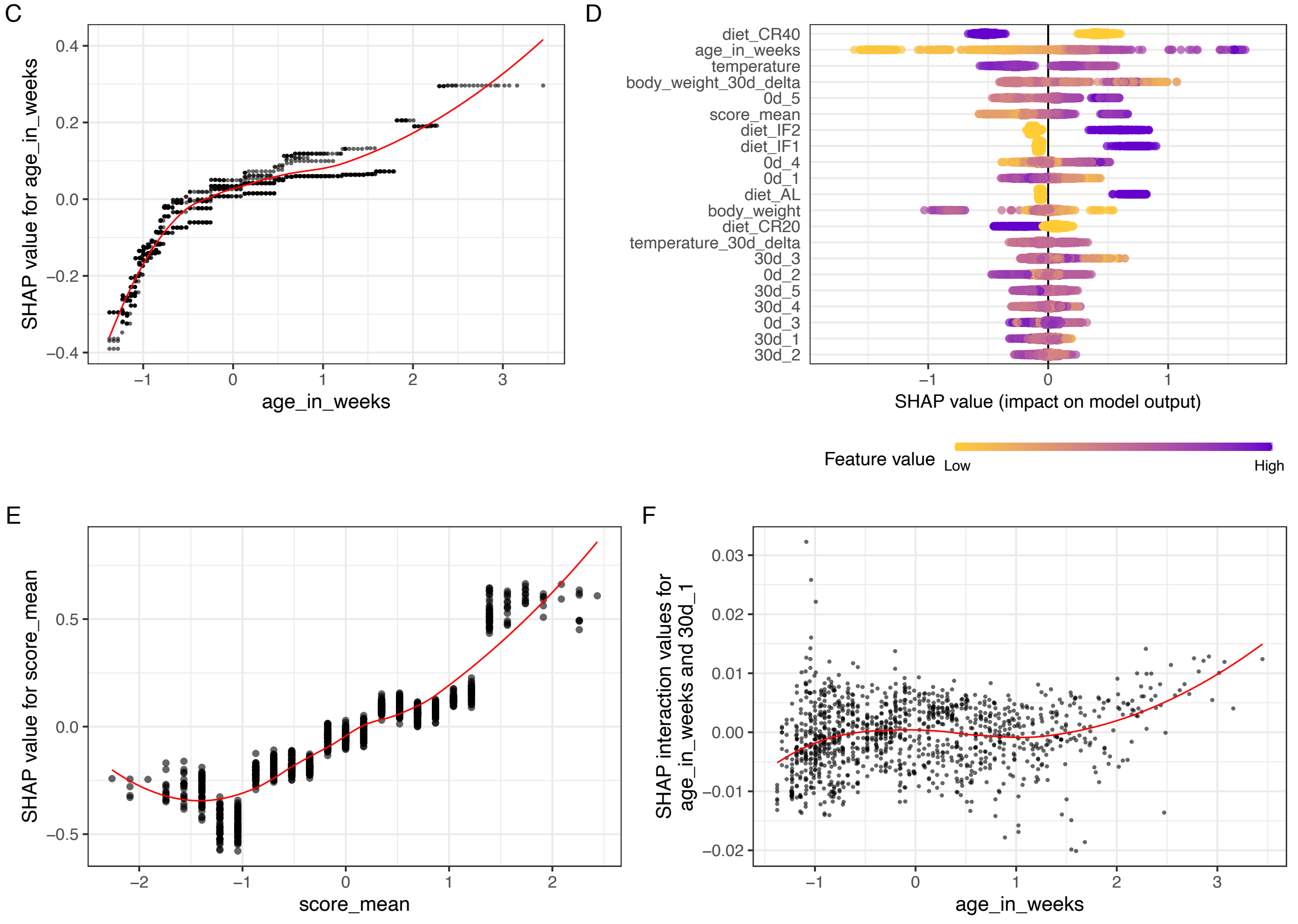

**eTable 1:** Fragility items.

| System and Item | Potential Deficits |
|---|---|
| **Digestive** | |
| Diarrhea | Soft feces |
| Malocclusions | Uneven and/or overgrown exposed teeth |
| Rectal Prolapse | Rectal tissue exposed |
| **Discomfort** | |
| Piloerection | Widespread piloerection |
| Head Piloerection[a] | Piloerection across the head |
| Response To Ocular Analgesic | Squinting unresolved |
| **Integument** | |
| Coat Condition | Coat is unkempt |
| Dermatitis | Skin lesions |
| Pallor and/or Cyanosis | Pale or cyanosis of skin |
| Dehydration/Skin Turgor | Tented skin return to normal $>$2 seconds |
| **Physical/Musculoskeletal** | |
| Distended Abdomen | Bulding of abdomen caudal to ribcage |
| Tumors | Visible or palpable mass |
| Thoracic Mass[a] | Visible or palpable mass |
| Body Condition | Body Condition Score 3 or less |
| Tremor | Tremor at rest |
| Kyphosis | Dorsal curvature of the spine |
| Hunched | Dorsal curvature of the spine with abdominal tuck |
| Gait Disorders | Hopping, wobbling, circling, wide stance or weakness |
| Tail Stiffening | Tail unresponsive when stroked |
| Paralysis | Paralysis of $>$1 limb |
| **Ocular/Nasal** | |
| Nasal Discharge | Nasal discharge, both nares |
| Eye Discharge or Swelling | Eye bulging and/or secretions |
| Changes to the Globe | Clouding and/or spotting of cornea or enlarged globe |
| Peri Retro Orbital Swelling | Swelling around eye $>$5mm |
| **Digestive/Urogenital** | |
| Urine | Bladder expression absent with massage |
| Vaginal/Uterine Prolapse | Vaginal/uterine tissue exposed |
| **Respiratory** | |
| Breathing Rate/Depth | Abnormal rate or effort, gasping |
| **Vestibulocochlear** | |
| Vestibular Disturbance | Head tilt, spinning, circling, head tuck, or trunk curling |
| **Behavior** | |
| Activity | No movement detected when placed in new cage |
| Response to External Stimuli | No response when lateral thorax nudged with a cotton tipped applicator |

[a]added to the index based on summary observations after the first six of 200 collection dates; rectal body temperature and body weight also measured; systems categories correspond to those in Whitehead *et al.* [13]

**eTable 2:** Cumulative incidence of frailty indicators by diet in aged J:DO mice

| | AL (N=25) | IF1 (N=33) | CR20 (N=55) | IF2 (N=48) | CR40 (N=73) | P-value |
|---|---|---|---|---|---|---|
| Activity | | | | | | |
| Absent | 14 (56.0%) | 24 (72.7%) | 28 (50.9%) | 31 (64.6%) | 41 (56.2%) | 0.0345 |
| Mild | 7 (28.0%) | 8 (24.2%) | 24 (43.6%) | 15 (31.3%) | 32 (43.8%) | |
| Severe | 4 (16.0%) | 1 (3.0%) | 3 (5.5%) | 2 (4.2%) | 0 (0%) | |
| Body Condition | | | | | | |
| Absent | 1 (4.0%) | 0 (0%) | 2 (3.6%) | 2 (4.2%) | 0 (0%) | <0.001 |
| Mild | 8 (32.0%) | 13 (39.4%) | 8 (14.5%) | 9 (18.8%) | 3 (4.1%) | |
| Severe | 16 (64.0%) | 20 (60.6%) | 45 (81.8%) | 37 (77.1%) | 70 (95.9%) | |
| Breathing Rate/Depth | | | | | | |
| Absent | 1 (4.0%) | 2 (6.1%) | 1 (1.8%) | 5 (10.4%) | 2 (2.7%) | 0.061 |
| Mild | 21 (84.0%) | 24 (72.7%) | 40 (72.7%) | 41 (85.4%) | 60 (82.2%) | |
| Severe | 3 (12.0%) | 7 (21.2%) | 14 (25.5%) | 2 (4.2%) | 11 (15.1%) | |
| Changes to Eye Globe | | | | | | |
| Absent | 10 (40.0%) | 19 (57.6%) | 20 (36.4%) | 24 (50.0%) | 19 (26.0%) | 0.0035 |
| Mild | 6 (24.0%) | 7 (21.2%) | 10 (18.2%) | 5 (10.4%) | 8 (11.0%) | |
| Severe | 9 (36.0%) | 7 (21.2%) | 25 (45.5%) | 19 (39.6%) | 46 (63.0%) | |
| Coat Condition | | | | | | |
| Absent | 0 (0%) | 0 (0%) | 0 (0%) | 0 (0%) | 0 (0%) | 0.012 |
| Mild | 3 (12.0%) | 0 (0%) | 4 (7.3%) | 3 (6.3%) | 0 (0%) | |
| Severe | 22 (88.0%) | 33 (100%) | 51 (92.7%) | 45 (93.8%) | 73 (100%) | |
| Dehydration, Skin Turgor | | | | | | |
| Absent | 2 (8.0%) | 7 (21.2%) | 6 (10.9%) | 8 (16.7%) | 3 (4.1%) | 0.058 |
| Mild | 0 (0%) | 0 (0%) | 0 (0%) | 0 (0%) | 0 (0%) | |
| Severe | 23 (92.0%) | 26 (78.8%) | 49 (89.1%) | 40 (83.3%) | 70 (95.9%) | |
| Dermatitis | | | | | | |
| Absent | 20 (80.0%) | 25 (75.8%) | 47 (85.5%) | 37 (77.1%) | 57 (78.1%) | 0.309 |
| Mild | 3 (12.0%) | 7 (21.2%) | 4 (7.3%) | 8 (16.7%) | 15 (20.5%) | |
| Severe | 2 (8.0%) | 1 (3.0%) | 4 (7.3%) | 3 (6.3%) | 1 (1.4%) | |
| Diarrhea | | | | | | |
| Absent | 24 (96.0%) | 32 (97.0%) | 55 (100%) | 47 (97.9%) | 73 (100%) | 0.161 |
| Severe | 1 (4.0%) | 1 (3.0%) | 0 (0%) | 1 (2.1%) | 0 (0%) | |
| Distended Abdomen | | | | | | |
| Absent | 0 (0%) | 1 (3.0%) | 5 (9.1%) | 3 (6.3%) | 21 (28.8%) | <0.001 |
| Mild | 4 (16.0%) | 5 (15.2%) | 8 (14.5%) | 16 (33.3%) | 23 (31.5%) | |
| Severe | 21 (84.0%) | 27 (81.8%) | 42 (76.4%) | 29 (60.4%) | 29 (39.7%) | |
| Eye Discharge/Eyelid Inflammation | | | | | | |
| Absent | 14 (56.0%) | 27 (81.8%) | 25 (45.5%) | 30 (62.5%) | 30 (41.1%) | <0.001 |
| Mild | 3 (12.0%) | 4 (12.1%) | 15 (27.3%) | 8 (16.7%) | 9 (12.3%) | |
| Severe | 8 (32.0%) | 2 (6.1%) | 15 (27.3%) | 10 (20.8%) | 34 (46.6%) | |
| Gait Disorders | | | | | | |
| Absent | 0 (0%) | 0 (0%) | 0 (0%) | 0 (0%) | 0 (0%) | 0.984 |
| Mild | 17 (68.0%) | 24 (72.7%) | 40 (72.7%) | 34 (70.8%) | 50 (68.5%) | |
| Severe | 8 (32.0%) | 9 (27.3%) | 15 (27.3%) | 14 (29.2%) | 23 (31.5%) | |
| Head Piloerection | | | | | | |
| Absent | 2 (8.3%) | 1 (3.1%) | 4 (7.5%) | 4 (9.3%) | 3 (4.3%) | 0.127 |
| Mild | 8 (33.3%) | 9 (28.1%) | 10 (18.9%) | 17 (39.5%) | 12 (17.1%) | |
| Severe | 14 (58.3%) | 22 (68.8%) | 39 (73.6%) | 22 (51.2%) | 55 (78.6%) | |
| Hunched | | | | | | |
| Absent | 4 (16.0%) | 9 (27.3%) | 1 (1.8%) | 6 (12.5%) | 1 (1.4%) | <0.001 |
| Mild | 7 (28.0%) | 7 (21.2%) | 8 (14.5%) | 6 (12.5%) | 4 (5.5%) | |
| Severe | 14 (56.0%) | 17 (51.5%) | 46 (83.6%) | 36 (75.0%) | 68 (93.2%) | |
| Kyphosis | | | | | | |
| Absent | 0 (0%) | 0 (0%) | 0 (0%) | 0 (0%) | 0 (0%) | 0.696 |
| Mild | 0 (0%) | 0 (0%) | 1 (1.8%) | 0 (0%) | 0 (0%) | |

| | | | | | | |
|---|---|---|---|---|---|---|
| Severe | 25 (100%) | 33 (100%) | 54 (98.2%) | 48 (100%) | 73 (100%) | |
| Malocclusions | | | | | | |
| Absent | 24 (96.0%) | 31 (93.9%) | 54 (98.2%) | 46 (95.8%) | 71 (97.3%) | 0.588 |
| Mild | 1 (4.0%) | 0 (0%) | 1 (1.8%) | 1 (2.1%) | 1 (1.4%) | |
| Severe | 0 (0%) | 2 (6.1%) | 0 (0%) | 1 (2.1%) | 1 (1.4%) | |
| Nasal Discharge | | | | | | |
| Absent | 25 (100%) | 33 (100%) | 55 (100%) | 48 (100%) | 73 (100%) | NA |
| Mild | 0 (0%) | 0 (0%) | 0 (0%) | 0 (0%) | 0 (0%) | |
| Severe | 0 (0%) | 0 (0%) | 0 (0%) | 0 (0%) | 0 (0%) | |
| Pallor/Cyanosis | | | | | | |
| Absent | 1 (4.0%) | 1 (3.0%) | 2 (3.6%) | 2 (4.2%) | 2 (2.7%) | 0.862 |
| Mild | 9 (36.0%) | 14 (42.4%) | 17 (30.9%) | 18 (37.5%) | 20 (27.4%) | |
| Severe | 15 (60.0%) | 18 (54.5%) | 36 (65.5%) | 28 (58.3%) | 51 (69.9%) | |
| Paralysis | | | | | | |
| Absent | 24 (96.0%) | 32 (97.0%) | 53 (96.4%) | 48 (100%) | 65 (89.0%) | 0.205 |
| Mild | 1 (4.0%) | 1 (3.0%) | 1 (1.8%) | 0 (0%) | 7 (9.6%) | |
| Severe | 0 (0%) | 0 (0%) | 1 (1.8%) | 0 (0%) | 1 (1.4%) | |
| Peri-retro-orbital Swelling | | | | | | |
| Absent | 23 (92.0%) | 32 (97.0%) | 43 (78.2%) | 43 (89.6%) | 58 (79.5%) | 0.152 |
| Mild | 1 (4.0%) | 1 (3.0%) | 8 (14.5%) | 4 (8.3%) | 6 (8.2%) | |
| Severe | 1 (4.0%) | 0 (0%) | 4 (7.3%) | 1 (2.1%) | 9 (12.3%) | |
| Piloerection | | | | | | |
| Absent | 0 (0%) | 0 (0%) | 0 (0%) | 0 (0%) | 0 (0%) | NA |
| Mild | 0 (0%) | 0 (0%) | 0 (0%) | 0 (0%) | 0 (0%) | |
| Severe | 25 (100%) | 33 (100%) | 55 (100%) | 48 (100%) | 73 (100%) | |
| Rectal Prolapse | | | | | | |
| Absent | 25 (100%) | 32 (97.0%) | 53 (96.4%) | 46 (95.8%) | 68 (93.2%) | 0.991 |
| Mild | 0 (0%) | 0 (0%) | 1 (1.8%) | 1 (2.1%) | 3 (4.1%) | |
| Severe | 0 (0%) | 1 (3.0%) | 1 (1.8%) | 1 (2.1%) | 2 (2.7%) | |
| Response to Analgesic | | | | | | |
| Absent | 21 (84.0%) | 30 (90.9%) | 49 (89.1%) | 40 (83.3%) | 66 (90.4%) | 0.593 |
| Mild | 4 (16.0%) | 3 (9.1%) | 3 (5.5%) | 6 (12.5%) | 6 (8.2%) | |
| Severe | 0 (0%) | 0 (0%) | 3 (5.5%) | 2 (4.2%) | 1 (1.4%) | |
| Response to External Stimuli | | | | | | |
| Absent | 25 (100%) | 33 (100%) | 55 (100%) | 48 (100%) | 72 (98.6%) | 1 |
| Severe | 0 (0%) | 0 (0%) | 0 (0%) | 0 (0%) | 1 (1.4%) | |
| Tail Stiffening | | | | | | |
| Absent | 4 (16.0%) | 6 (18.2%) | 6 (10.9%) | 4 (8.3%) | 5 (6.8%) | 0.807 |
| Mild | 8 (32.0%) | 10 (30.3%) | 17 (30.9%) | 14 (29.2%) | 22 (30.1%) | |
| Severe | 13 (52.0%) | 17 (51.5%) | 32 (58.2%) | 30 (62.5%) | 46 (63.0%) | |
| Thoracic Mass | | | | | | |
| Absent | 12 (50.0%) | 19 (59.4%) | 19 (35.8%) | 23 (53.5%) | 27 (38.6%) | 0.481 |
| Mild | 8 (33.3%) | 7 (21.9%) | 20 (37.7%) | 13 (30.2%) | 26 (37.1%) | |
| Severe | 4 (16.7%) | 6 (18.8%) | 14 (26.4%) | 7 (16.3%) | 17 (24.3%) | |
| Tremor | | | | | | |
| Absent | 0 (0%) | 2 (6.1%) | 0 (0%) | 2 (4.2%) | 1 (1.4%) | 0.156 |
| Mild | 8 (32.0%) | 10 (30.3%) | 11 (20.0%) | 13 (27.1%) | 11 (15.1%) | |
| Severe | 17 (68.0%) | 21 (63.6%) | 44 (80.0%) | 33 (68.8%) | 61 (83.6%) | |
| Tumors | | | | | | |
| Absent | 4 (16.0%) | 13 (39.4%) | 9 (16.4%) | 18 (37.5%) | 26 (35.6%) | 0.0755 |
| Mild | 3 (12.0%) | 6 (18.2%) | 11 (20.0%) | 9 (18.8%) | 16 (21.9%) | |
| Severe | 18 (72.0%) | 14 (42.4%) | 35 (63.6%) | 21 (43.8%) | 31 (42.5%) | |
| Urine | | | | | | |
| Absent | 2 (8.0%) | 7 (21.2%) | 5 (9.1%) | 8 (16.7%) | 2 (2.7%) | 0.018 |
| Mild | 0 (0%) | 0 (0%) | 0 (0%) | 0 (0%) | 0 (0%) | |
| Severe | 23 (92.0%) | 26 (78.8%) | 50 (90.9%) | 40 (83.3%) | 71 (97.3%) | |
| Vaginal/Uterine Prolapse | | | | | | |
| Absent | 25 (100%) | 33 (100%) | 55 (100%) | 48 (100%) | 73 (100%) | NA |

| | | | | | | |
|---|---|---|---|---|---|---|
| Mild | 0 (0%) | 0 (0%) | 0 (0%) | 0 (0%) | 0 (0%) | |
| Severe | 0 (0%) | 0 (0%) | 0 (0%) | 0 (0%) | 0 (0%) | |
| Vestibular Disturbance | | | | | | |
| Absent | 4 (16.0%) | 4 (12.1%) | 3 (5.5%) | 10 (20.8%) | 1 (1.4%) | <0.001 |
| Mild | 11 (44.0%) | 18 (54.5%) | 26 (47.3%) | 18 (37.5%) | 17 (23.3%) | |
| Severe | 10 (40.0%) | 11 (33.3%) | 26 (47.3%) | 20 (41.7%) | 55 (75.3%) | |

**eTable 3:** Cumulative incidence of frailty indicators by sex in aged C57BL/6J mice

| | Males (N=24) | Females (N=12) | P-value |
|---|---|---|---|
| Activity | | | |
| Absent | 9 (37.5%) | 12 (100%) | <0.001 |
| Mild | 15 (62.5%) | 0 (0%) | |
| Severe | 0 (0%) | 0 (0%) | |
| Body Condition | | | |
| Absent | 1 (4.2%) | 0 (0%) | 0.82 |
| Mild | 10 (41.7%) | 6 (50.0%) | |
| Severe | 13 (54.2%) | 6 (50.0%) | |
| Breathing Rate/Depth | | | |
| Absent | 0 (0%) | 1 (8.3%) | 0.235 |
| Mild | 23 (95.8%) | 10 (83.3%) | |
| Severe | 1 (4.2%) | 1 (8.3%) | |
| Changes to Eye Globe | | | |
| Absent | 23 (95.8%) | 10 (83.3%) | 0.256 |
| Mild | 1 (4.2%) | 1 (8.3%) | |
| Severe | 0 (0%) | 1 (8.3%) | |
| Coat Condition | | | |
| Absent | 0 (0%) | 1 (8.3%) | 0.1 |
| Mild | 1 (4.2%) | 2 (16.7%) | |
| Severe | 23 (95.8%) | 9 (75.0%) | |
| Dehydration, Skin Turgor | | | |
| Absent | 11 (45.8%) | 7 (58.3%) | 0.727 |
| Mild | 0 (0%) | 0 (0%) | |
| Severe | 13 (54.2%) | 5 (41.7%) | |
| Dermatitis | | | |
| Absent | 19 (79.2%) | 12 (100%) | 0.367 |
| Mild | 4 (16.7%) | 0 (0%) | |
| Severe | 1 (4.2%) | 0 (0%) | |
| Diarrhea | | | |
| Absent | 23 (95.8%) | 12 (100%) | 1 |
| Severe | 1 (4.2%) | 0 (0%) | |
| Distended Abdomen | | | |
| Absent | 1 (4.2%) | 1 (8.3%) | 0.336 |
| Mild | 3 (12.5%) | 3 (25.0%) | |
| Severe | 20 (83.3%) | 8 (66.7%) | |
| Eye Discharge/Eyelid Inflammation | | | |
| Absent | 18 (75.0%) | 6 (50.0%) | 0.174 |
| Mild | 6 (25.0%) | 5 (41.7%) | |
| Severe | 0 (0%) | 1 (8.3%) | |
| Gait Disorders | | | |
| Absent | 0 (0%) | 0 (0%) | 1 |
| Mild | 22 (91.7%) | 11 (91.7%) | |
| Severe | 2 (8.3%) | 1 (8.3%) | |
| Head Piloerection | | | |
| Absent | 1 (4.2%) | 4 (33.3%) | 0.067 |
| Mild | 11 (45.8%) | 3 (25.0%) | |
| Severe | 12 (50.0%) | 5 (41.7%) | |
| Hunched | | | |
| Absent | 0 (0%) | 0 (0%) | 0.028 |
| Mild | 2 (8.3%) | 5 (41.7%) | |
| Severe | 22 (91.7%) | 7 (58.3%) | |
| Kyphosis | | | |
| Absent | 0 (0%) | 0 (0%) | 0.342 |
| Mild | 0 (0%) | 1 (8.3%) | |

| | | | |
|---|---|---|---|
| Severe | 24 (100%) | 11 (91.7%) | |
| Malocclusions | | | |
| Absent | 24 (100%) | 12 (100%) | NA |
| Mild | 0 (0%) | 0 (0%) | |
| Severe | 0 (0%) | 0 (0%) | |
| Nasal Discharge | | | |
| Absent | 24 (100%) | 12 (100%) | NA |
| Mild | 0 (0%) | 0 (0%) | |
| Severe | 0 (0%) | 0 (0%) | |
| Pallor/Cyanosis | | | |
| Absent | 1 (4.2%) | 2 (16.7%) | 0.499 |
| Mild | 15 (62.5%) | 6 (50.0%) | |
| Severe | 8 (33.3%) | 4 (33.3%) | |
| Paralysis | | | |
| Absent | 24 (100%) | 12 (100%) | NA |
| Mild | 0 (0%) | 0 (0%) | |
| Severe | 0 (0%) | 0 (0%) | |
| Peri-retro-orbital Swelling | | | |
| Absent | 23 (95.8%) | 9 (75.0%) | 0.101 |
| Mild | 1 (4.2%) | 3 (25.0%) | |
| Severe | 0 (0%) | 0 (0%) | |
| Piloerection | | | |
| Absent | 0 (0%) | 0 (0%) | NA |
| Mild | 0 (0%) | 0 (0%) | |
| Severe | 24 (100%) | 12 (100%) | |
| Rectal Prolapse | | | |
| Absent | 24 (100%) | 10 (83.3%) | 0.109 |
| Mild | 0 (0%) | 0 (0%) | |
| Severe | 0 (0%) | 2 (16.7%) | |
| Response to Analgesic | | | |
| Absent | 24 (100%) | 12 (100%) | NA |
| Mild | 0 (0%) | 0 (0%) | |
| Severe | 0 (0%) | 0 (0%) | |
| Response to External Stimuli | | | |
| Absent | 24 (100%) | 12 (100%) | 1 |
| Severe | 0 (0%) | 0 (0%) | |
| Tail Stiffening | | | |
| Absent | 1 (4.2%) | 2 (16.7%) | 0.147 |
| Mild | 10 (41.7%) | 2 (16.7%) | |
| Severe | 13 (54.2%) | 8 (66.7%) | |
| Thoracic Mass | | | |
| Absent | 11 (45.8%) | 9 (75.0%) | 0.158 |
| Mild | 13 (54.2%) | 3 (25.0%) | |
| Severe | 0 (0%) | 0 (0%) | |
| Tremor | | | |
| Absent | 0 (0%) | 1 (8.3%) | 0.516 |
| Mild | 10 (41.7%) | 4 (33.3%) | |
| Severe | 14 (58.3%) | 7 (58.3%) | |
| Tumors | | | |
| Absent | 2 (8.3%) | 2 (16.7%) | 0.0035 |
| Mild | 2 (8.3%) | 6 (50.0%) | |
| Severe | 20 (83.3%) | 4 (33.3%) | |
| Urine | | | |
| Absent | 11 (45.8%) | 7 (58.3%) | 0.727 |
| Mild | 0 (0%) | 0 (0%) | |
| Severe | 13 (54.2%) | 5 (41.7%) | |
| Vaginal/Uterine Prolapse | | | |
| Absent | 24 (100%) | 12 (100%) | NA |

| | | | |
|---|---|---|---|
| Mild | 0 (0%) | 0 (0%) | |
| Severe | 0 (0%) | 0 (0%) | |
| Vestibular Disturbance | | | |
| Absent | 0 (0%) | 2 (16.7%) | 0.139 |
| Mild | 5 (20.8%) | 3 (25.0%) | |
| Severe | 19 (79.2%) | 7 (58.3%) | |

**eTable 4:** Correlation coefficients (95% confidence intervals [CI]) and p-values for life expectancy with fragility items by diet in aged J:DO mice

| | r AL | p AL | r IF1 | p IF1 | r CR20 | p CR20 | r IF2 | p IF2 | r CR40 | p CR40 |
|---|---|---|---|---|---|---|---|---|---|---|
| Body Condition | -0.40 | <0.001 | -0.34 | <0.001 | -0.33 | <0.001 | -0.36 | <0.001 | -0.26 | <0.001 |
| Body Weight | 0.33 | <0.001 | 0.35 | <0.001 | 0.14 | <0.001 | 0.42 | <0.001 | 0.28 | <0.001 |
| Tail Stiffening | -0.30 | <0.001 | -0.24 | <0.001 | -0.10 | 0.001 | -0.25 | <0.001 | -0.28 | <0.001 |
| Head Piloerection | -0.29 | <0.001 | -0.21 | <0.001 | -0.24 | <0.001 | -0.35 | <0.001 | -0.31 | <0.001 |
| Gait Disorders | -0.29 | <0.001 | -0.24 | <0.001 | -0.10 | 0.001 | -0.08 | 0.042 | -0.12 | <0.001 |
| Tremor | -0.28 | <0.001 | -0.16 | 0.001 | -0.26 | <0.001 | -0.15 | <0.001 | -0.27 | <0.001 |
| Temperature | 0.27 | <0.001 | 0.26 | <0.001 | 0.24 | <0.001 | 0.30 | <0.001 | 0.27 | <0.001 |
| FgI Score | -0.26 | <0.001 | -0.26 | <0.001 | -0.37 | <0.001 | -0.33 | <0.001 | -0.37 | <0.001 |
| Eye Discharge/Eyelid Inflammation | 0.24 | <0.001 | 0.17 | <0.001 | -0.18 | <0.001 | -0.05 | 0.23 | -0.14 | <0.001 |
| Tumors | -0.23 | <0.001 | -0.12 | 0.014 | -0.26 | <0.001 | -0.24 | <0.001 | -0.14 | <0.001 |
| Age in weeks | -0.21 | <0.001 | -0.35 | <0.001 | -0.44 | <0.001 | -0.37 | <0.001 | -0.43 | <0.001 |
| Coat Condition | -0.20 | <0.001 | -0.17 | 0.001 | -0.29 | <0.001 | -0.18 | <0.001 | -0.23 | <0.001 |
| Breathing Rate/Depth | -0.19 | <0.001 | -0.25 | <0.001 | -0.29 | <0.001 | -0.23 | <0.001 | -0.19 | <0.001 |
| N Deficits | -0.19 | <0.001 | -0.21 | <0.001 | -0.34 | <0.001 | -0.29 | <0.001 | -0.35 | <0.001 |
| Changes to Eye Globe | 0.18 | 0.001 | 0.28 | <0.001 | -0.04 | 0.14 | -0.11 | 0.005 | -0.12 | <0.001 |
| Kyphosis | -0.16 | 0.003 | -0.15 | 0.002 | -0.08 | 0.004 | -0.09 | 0.02 | -0.10 | <0.001 |
| Hunched | -0.16 | 0.003 | -0.35 | <0.001 | -0.18 | <0.001 | -0.22 | <0.001 | -0.23 | <0.001 |
| Thoracic Mass | -0.16 | 0.003 | -0.10 | 0.036 | -0.14 | <0.001 | -0.13 | 0.001 | -0.05 | 0.012 |
| Dermatitis | -0.15 | 0.004 | -0.12 | 0.013 | -0.06 | 0.029 | -0.18 | <0.001 | -0.08 | <0.001 |
| Dehydration, Skin Turgor | -0.14 | 0.012 | -0.03 | 0.53 | -0.12 | <0.001 | -0.14 | <0.001 | -0.14 | <0.001 |
| Pallor/Cyanosis | -0.13 | 0.016 | -0.10 | 0.041 | -0.11 | <0.001 | -0.21 | <0.001 | -0.21 | <0.001 |
| Distended Abdomen | 0.10 | 0.053 | 0.03 | 0.6 | 0.07 | 0.012 | 0.18 | <0.001 | 0.03 | 0.19 |
| Malocclusions | -0.06 | 0.25 | -0.04 | 0.47 | -0.03 | 0.34 | -0.01 | 0.7 | 0.01 | 0.78 |
| Diarrhea | -0.04 | 0.44 | -0.05 | 0.31 | | | -0.04 | 0.25 | | |
| Response to Analgesic | -0.02 | 0.67 | -0.07 | 0.17 | -0.06 | 0.047 | 0.13 | 0.001 | -0.06 | 0.003 |
| Peri-retro-orbital Swelling | -0.02 | 0.68 | -0.06 | 0.21 | -0.08 | 0.006 | -0.12 | 0.002 | -0.05 | 0.015 |
| Piloerection | -0.01 | 0.79 | -0.05 | 0.33 | -0.08 | 0.008 | -0.14 | <0.001 | -0.04 | 0.037 |
| Activity | 0.01 | 0.77 | 0.01 | 0.9 | 0.00 | 0.93 | -0.04 | 0.28 | -0.03 | 0.12 |
| Paralysis | -0.01 | 0.86 | -0.03 | 0.5 | -0.06 | 0.029 | | | -0.11 | <0.001 |
| Vestibular Disturbance | -0.01 | 0.89 | 0.00 | 0.94 | -0.05 | 0.09 | -0.01 | 0.75 | -0.07 | <0.001 |
| Urine | 0.01 | 0.92 | -0.01 | 0.77 | -0.10 | <0.001 | -0.11 | 0.004 | -0.12 | <0.001 |
| Nasal Discharge | | | | | | | | | | |
| Rectal Prolapse | | | -0.05 | 0.31 | -0.05 | 0.11 | -0.09 | 0.019 | -0.07 | <0.001 |
| Vaginal/Uterine Prolapse | | | | | | | | | | |
| Response to External Stimuli | | | | | | | | | -0.03 | 0.18 |

Items ordered by descending absolute correlation coefficient in AL cohort; r=Pearson correlation.

**eTable 5:** Correlation coefficients (95% confidence intervals [CI]) and p-values for life expectancy with fragility items by sex in aged C57BL/6J mice

| | **r Females** | **p Females** | **r Males** | **p Males** |
|---|---|---|---|---|
| Age in weeks | -0.45 | <0.001 | -0.47 | <0.001 |
| FgI Score | -0.44 | <0.001 | -0.42 | <0.001 |
| Body Condition | -0.44 | <0.001 | -0.37 | <0.001 |
| Pallor/Cyanosis | -0.40 | <0.001 | -0.23 | <0.001 |
| Head Piloerection | -0.39 | <0.001 | -0.39 | <0.001 |
| N Deficits | -0.34 | <0.001 | -0.32 | <0.001 |
| Coat Condition | -0.34 | <0.001 | -0.14 | <0.001 |
| Body Weight | 0.33 | <0.001 | 0.09 | 0.02 |
| Hunched | -0.33 | <0.001 | -0.29 | <0.001 |
| Tremor | -0.33 | <0.001 | -0.30 | <0.001 |
| Tumors | -0.26 | <0.001 | -0.36 | <0.001 |
| Eye Discharge/Eyelid Inflammation | -0.25 | <0.001 | -0.12 | <0.001 |
| Temperature | 0.24 | <0.001 | 0.28 | <0.001 |
| Rectal Prolapse | -0.22 | <0.001 | | |
| Urine | -0.21 | <0.001 | -0.09 | 0.03 |
| Dehydration, Skin Turgor | -0.21 | <0.001 | -0.13 | <0.001 |
| Tail Stiffening | -0.18 | 0.01 | -0.17 | <0.001 |
| Breathing Rate/Depth | -0.17 | 0.01 | -0.19 | <0.001 |
| Distended Abdomen | 0.17 | 0.01 | -0.21 | <0.001 |
| Changes to Eye Globe | -0.16 | 0.01 | 0.03 | 0.37 |
| Peri-retro-orbital Swelling | -0.15 | 0.03 | 0.02 | 0.54 |
| Kyphosis | -0.14 | 0.04 | -0.10 | 0.01 |
| Vestibular Disturbance | 0.13 | 0.04 | 0.06 | 0.13 |
| Gait Disorders | -0.07 | 0.32 | -0.18 | <0.001 |
| Thoracic Mass | 0.05 | 0.46 | -0.05 | 0.21 |
| Piloerection | -0.02 | 0.76 | 0.06 | 0.1 |
| Dermatitis | | | -0.13 | <0.001 |
| Response to Analgesic | | | | |
| Nasal Discharge | | | | |
| Vaginal/Uterine Prolapse | | | | |
| Diarrhea | | | 0.03 | 0.48 |
| Activity | | | 0.04 | 0.31 |
| Response to External Stimuli | | | | |
| Paralysis | | | | |
| Malocclusions | | | | |

Items ordered by descending absolute correlation coefficient in Female cohort; r=Pearson correlation.

**eTable 6:** Model performance for TxW predictive endpoint criteria across two genetic backgrounds, both sexes, and five experimental groups.

| Strain | Sex | Diet | Predicted | Observed | Sensitivity | Specificity | NPV | PPV |
|---|---|---|---|---|---|---|---|---|
| C57BL/6J | M | AL | 85 | 109 | 0.28 | 0.84 | 0.79 | 0.36 |
| C57BL/6J | F | AL | 82 | 53 | 0.93 | 0.48 | 0.89 | 0.60 |
| J:DO | F | IF1 | 85 | 116 | 0.53 | 0.61 | 0.40 | 0.73 |
| J:DO | F | CR20 | 280 | 290 | 0.45 | 0.67 | 0.66 | 0.46 |
| J:DO | F | IF2 | 130 | 174 | 0.53 | 0.74 | 0.56 | 0.71 |
| J:DO | F | CR40 | 391 | 434 | 0.26 | 0.80 | 0.77 | 0.29 |
| J:DO | F | AL | 55 | 70 | 0.27 | 0.46 | 0.38 | 0.34 |

Note: Abbreviations: NPV = negative predicted value, PPV = positive predicted value, TxW = the product of temperature and body weight. PPV is the probability that, following a positive prediction, the sample is truly positive. NPV is the probability that, following a negative prediction, the sample is truly negative. TxW performance required 13+ weeks of data.
